# The Genetic Architecture of DNA Replication Timing in Human Pluripotent Stem Cells

**DOI:** 10.1101/2020.05.08.085324

**Authors:** Qiliang Ding, Matthew M. Edwards, Michelle L. Hulke, Alexa N. Bracci, Ya Hu, Yao Tong, Xiang Zhu, Joyce Hsiao, Christine J. Charvet, Sulagna Ghosh, Robert E. Handsaker, Kevin Eggan, Florian T. Merkle, Jeannine Gerhardt, Dieter Egli, Andrew G. Clark, Amnon Koren

## Abstract

DNA replication follows a strict spatiotemporal program that intersects with chromatin structure and gene regulation. However, the genetic basis of the mammalian DNA replication timing program is poorly understood^1–3^. To systematically identify genetic regulators of DNA replication timing, we exploited inter-individual variation in 457 human pluripotent stem cell lines from 349 individuals. We show that the human genome’s replication program is broadly encoded in DNA and identify 1,617 *cis*-acting replication timing quantitative trait loci (rtQTLs^4^) – base-pair-resolution sequence determinants of replication initiation. rtQTLs function individually, or in combinations of proximal and distal regulators, to affect replication timing. Analysis of rtQTL locations reveals a histone code for replication initiation, composed of bivalent histone H3 trimethylation marks on a background of histone hyperacetylation. The H3 trimethylation marks are individually repressive yet synergize to promote early replication. We further identify novel positive and negative regulators of DNA replication timing, the former comprised of pluripotency-related transcription factors while the latter involve boundary elements. Human replication timing is controlled by a multi-layered mechanism that operates on target DNA sequences, is composed of dozens of effectors working combinatorially, and follows principles analogous to transcription regulation: a histone code, activators and repressors, and a promoter-enhancer logic.

## Main

Eukaryotic genomes are replicated according to a strict spatiotemporal program, in which replication initiates from specific locations along chromosomes and at reproducible times. The replication timing program is a fundamental property of chromosome organization, interfaces with gene regulation and shapes the mutational landscape of the genome. Efforts to understand the locations and nature of initiation sites and the factors that regulate DNA replication timing in mammalian cells have been ongoing for decades, with limited success^1–3^. Specifically, it is still unclear to what extent the DNA replication timing program is determined by local DNA sequences, by epigenetic factors, or by a combination thereof. Earlier studies suggested that specific sequence elements control replication initiation in human cells, with several distal and proximal elements often acting in concert^5–11^. More recently, CRISPR/Cas9-mediated deletions have suggested that several DNA sequences locally interact to control early replication in mice^12^.

Numerous lines of evidence link replication regulation to epigenetic states, in particular histone acetylations and methylations marking open chromatin^3,13–17^. However, no single epigenetic mark appears to be absolutely required nor sufficient for replication origin function. This has led to suggestions that a combination of histone marks may be required for specifying patterns of DNA replication^18^. Similarly, it has been proposed that indiscriminate DNA-binding patterns of the replication machinery may translate into a consistent, organized replication program by means of combinatorial chromatin modifications influencing subsets of replication initiation sites^3^.

The nature of such modular, combinatorial regulation of DNA replication at the genetic and epigenetic levels remains to be revealed. Previous studies applied stepwise reverse engineering approaches to probe for mechanisms controlling replication timing. However, such a complex system may be best studied with an unbiased and comprehensive interrogation of genetic and epigenetic factors and their interactions. While such an approach is currently challenging experimentally, an alternative is to take advantage of natural genetic variation. We previously showed that replication timing is variable among individuals, that it can be studied at fine-scale on a population level by sequencing the genomes of proliferating cells, and that genotype information from the same genome sequences can be used to associate replication timing variation with specific genetic polymorphisms. This results in the identification of replication timing quantitative trait loci (rtQTLs), DNA sequences that act in *cis* to affect replication initiation^4^. Leveraging human genetic variation enables the equivalent of numerous surgical genetic manipulations and their association with DNA replication timing alterations.

Here, we apply this approach to hundreds of human embryonic stem cell (hESC) and induced pluripotent stem cell (iPSC) lines. Pluripotent stem cells are particularly useful for this analysis, since they are non-transformed, karyotypically stable and highly proliferative, and have a wealth of epigenetic data available for multi-omic analyses. We identify 1,617 *cis*-rtQTLs and analyze their locations and allelic differences. These analyses delineate the architecture of human replication timing as a quantitative trait involving combinatorial regulation by several layers of epigenetic mechanisms rooted in *cis*-acting DNA sequences.

### High-resolution population-scale replication timing profiles

To comprehensively characterize human inter-individual replication timing variation and its genetic basis, we analyzed deep (∼30x) whole-genome sequences of 121 hESC lines and 326 iPSC lines^19^ and sequenced another 24 hESCs and 17 iPSCs for a total of 488 cell lines (Methods). ES and iPS cultures are highly proliferative, containing 35–55% cells in S phase.

DNA replication timing leads to variation in DNA copy number along chromosomes among S phase cells (e.g., early-replicating regions are duplicated in most cells), causing read depth fluctuations in the sequencing data^4^. Indeed, we were able to generate high-resolution replication timing profiles for a total of 140 hESCs and 317 iPSCs (Methods). ES and iPS cells had similar replication profiles, as expected.

Replication timing profiles were continuous along chromosomes, highly reproducible among samples (median *r* = 0.93), and consistent with previous replication timing measurements by Repli-Seq (median *r* = 0.86; Fig. 1, A–D). The replication profiles were exceptionally sharp, in line with recent high-resolution Repli-Seq data^20^, with discrete peaks and valleys (local maxima and minima) that were themselves highly reproducible among individuals. Replication timing peaks represent prominent initiation sites containing one or more replication origins. We further improved data resolution using principal component (PC)-based correction across cell lines (Fig. 1, C and D; Methods).

**Figure 1.**
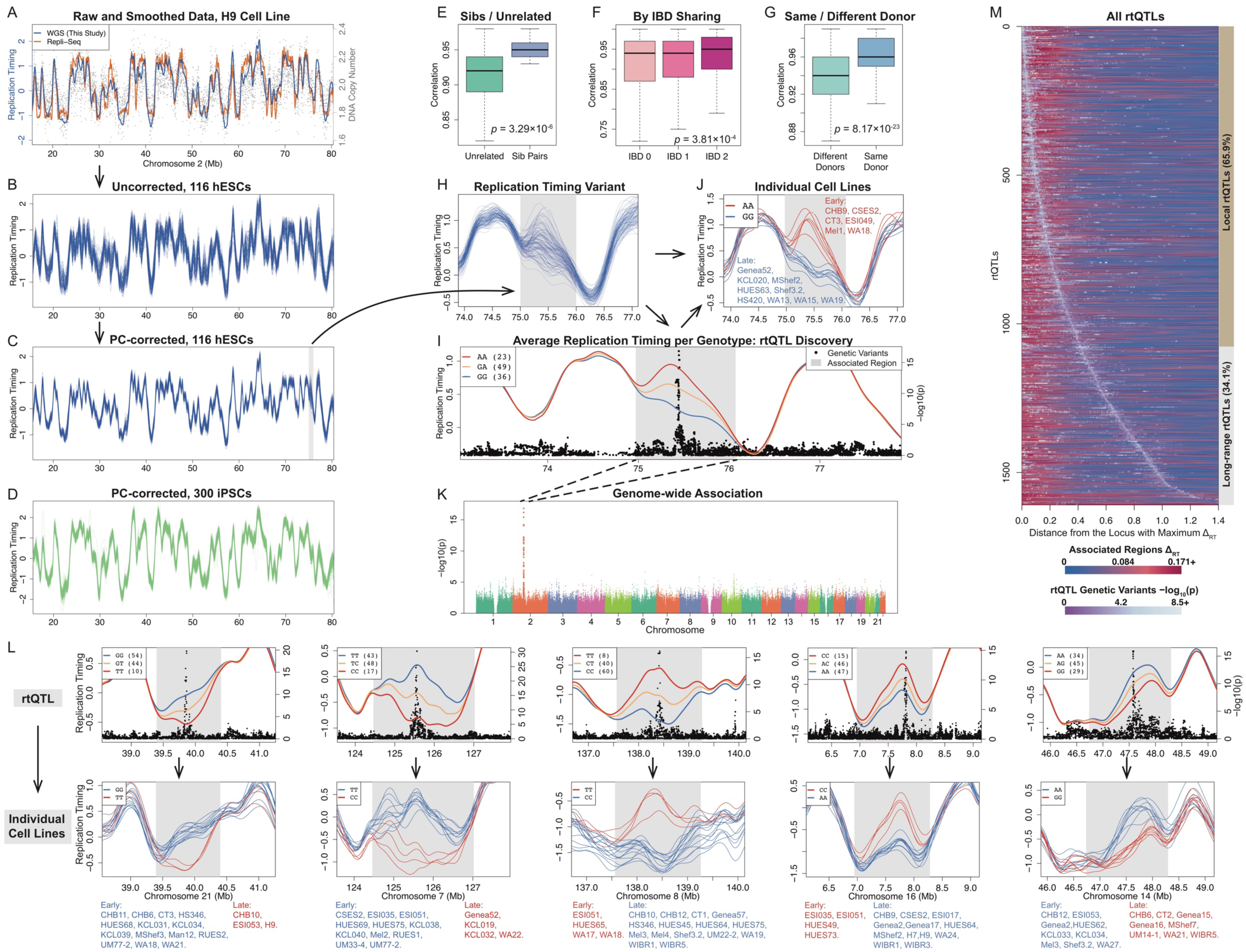
The Human Genome’s Replication Timing Program is Extensively Encoded in DNA. (A) Replication timing inferred from read depth fluctuations. Read depth (gray dots) and replication timing profile (blue line; *Z*-score) of the H9 cell line. Green line: Repli-Seq data of the same cell line.^23^ (B) Replication timing profiles are highly reproducible among samples. (C, D) Leveraging the population-scale of the data, PC-based correction greatly improves replication profile accuracy. (E–G) Genetic relatedness is associated with replication timing similarity. (E) Comparison of sibling vs. unrelated hESC lines. (F) Genomic regions stratified by increasing identity-by-descent. (G) iPSCs from the same or different donors. (H) A genomic region (gray) with substantial inter-individual replication timing variation. (I–K) Genetic association reveals rtQTLs. (I) A SNP haplotype strongly associates in *cis* with the replication timing variant from panel H (panel K shows the genome-wide association). Mean replication timing profiles (left Y axis) for individuals with different genotypes at rs12713840, the top SNP, demonstrates that SNPs in *cis* (right Y axis) associate with replication initiation activity. Gray shaded area represents the affected genomic region. (J) Replication timing at the variant from panel H, stratifying individuals by rs12713840 genotype, demonstrates that genotype is the main determinant of replication timing variation. (L) Additional rtQTL examples. Similar to panels I and J. Most rtQTLs affect peaks (replication initiation regions). (M) All rtQTLs. Each horizontal line is an rtQTL, oriented from the replication timing locus with maximum difference between early-and late-replicating genotypes (*Δ*_*RT*_) and showing the averaged replication timing difference on both sides of that locus (i.e., the rtQTL-associated region spans twice the distance shown; refer to panel I). Foreground (gray-purple) shades are the rtQTL SNPs, color-coded by *p*-values, and placed according to their distance to the locus of maximal *Δ*_*RT*_. rtQTLs are encoded in localized haplotypes yet influence extended genomic regions up to 5.6 Mb. Most rtQTLs influence surrounding genomic region (“local”), while a subset show long-range effects.

### DNA replication timing is broadly influenced by *cis*-acting sequences

While replication timing profiles were highly reproducible among individuals, we nonetheless observed genomic regions with substantial inter-individual variation. We identified 1,489 autosomal replication timing variants in hESCs and 1,837 in iPSCs, cumulatively encompassing 795 Mb (34%) and 980 Mb (40.8%), respectively, of the analyzable genome (Fig. 1, C and D). We hypothesized that at least some of this variation is due to genetic polymorphism. To test this, we first compared replication timing variation between 24 pairs of hESC lines that are genetic siblings, versus unrelated cell lines; between genomic regions that are identical by descent (IBD), half-identical or non-identical between sibling cell lines; and between 108 pairs of iPSC lines derived from the same donor, compared to different donors (Methods). Consistent with a significant genetic contribution to replication timing variation, samples or genomic intervals that are genetically related consistently showed greater replication timing similarity than unrelated comparisons (Fig. 1, E–G).

To further dissect genetic contributions to replication timing variation, we used our previously described rtQTL mapping approach^4^ to associate replication timing with specific genetic polymorphisms. This approach was applied here at larger scale, to deeper-sequenced data, and with refined algorithms than before (Methods). We limited this analysis to 108 hESCs of European ancestry and to 192 iPSCs from different individuals.

We identified 1,617 *cis*-rtQTLs (FDR 0.1; 1,012 were identified with FDR 0.05; Fig. 1, I–M; Table S1), two orders of magnitude more than previous associations of replication timing with *cis*-acting sequences^4,12^. We used CAVIAR^21^ to fine-map (90% credible level) a median of 33 SNPs per rtQTL, with 316 rtQTLs mapped to within 10 SNPs and 36 rtQTLs mapped to no more than three SNPs. rtQTL mapping was cross-validated between ES and iPS cells and further confirmed using additionally-sequenced cell lines and with a locus-specific single-molecule assay (Fig. S1).

rtQTLs influenced the replication timing of regions spanning 858 kb on average and a total of 741.8 Mb of genomic sequence (31.8% of the genome, Fig. 1M). This is a lower bound estimate of the extent to which human replication timing is influenced by DNA sequence, since our approach will not detect weaker rtQTLs, invariant sequences or rare variant rtQTLs. Intriguingly, 67.9% (1,098) of rtQTLs coincided with sharp peaks in the replication profiles (binomial *p* = 2.24×10^−25^; Fig. 1, I–L), and rtQTL SNPs were significantly closer to peaks than expected (Wilcoxon rank-sum *p* = 1.77×10^−16^). This suggests that rtQTLs may influence replication initiation, as previously reported^4,22^, and that most rtQTLs can be used as fine-scale markers of replication initiation regions. The identification of rtQTLs as precise genetic determinants of replication timing provides a unique opportunity to fine-map molecular mechanisms controlling replication initiation and timing.

**Figure S1.**
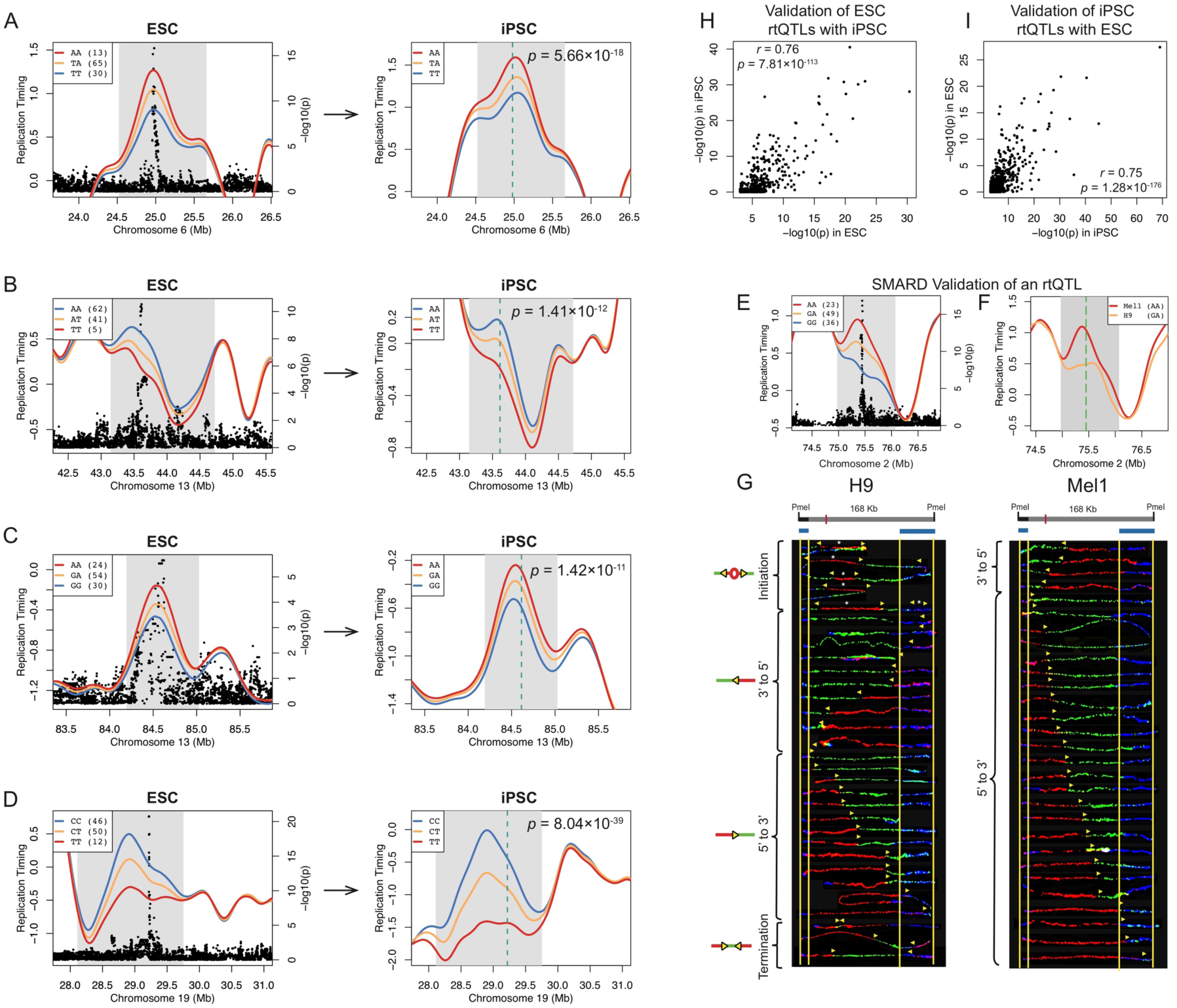
rtQTL Validation. (A–D) Validation of rtQTLs in 192 iPSC lines (Methods). The left panels are examples of rtQTLs in hESCs. The right panels show replication timing in the same regions in iPSCs, stratified by the genotype of the top rtQTL SNP discovered in the hESCs (vertical line). Association *p*-values in iPSCs are indicated. Excellent agreement between hESCs and iPSCs demonstrate that the rtQTLs discovered in hESCs are reproducible in an independent cohort. (E–G) SMARD (single-molecule analysis of replicated DNA ^22^) analysis of an rtQTL on chromosome 2 (Fig. 1I) in Mel1 and H9 cell lines confirms variation in initiation site activity consistent with rtQTL genotypes. (F) Replication timing flanking the rtQTL locus (gray region); green line: the region analyzed by SMARD. The initiation site on the left side of the green line is an rtQTL (panel E), at which Mel1 and H9 carry the early-replicating and heterozygous genotype, respectively. (G) SMARD results, where each line indicates one DNA molecule, and the shift from red to green reveals the location and direction of replication forks (yellow arrows). Significantly more forks are progressing from 5’ to 3’ in Mel1 when compared with H9 (*p* = 0.027, Fisher’s exact test), indicating that the upstream initiation site is much stronger in Mel1 than H9, consistent with the rtQTL analysis. (H, I) rtQTLs are highly reproducible between the ESCs and iPSCs. When directly testing ESC rtQTLs using iPSCs (H) or *vice versa* (I), the *p*-values show strong positive correlation. Among the 602 ESC rtQTLs tested, 38.7% (233/602) were validated (*p* < 0.05 and the same direction of effect) in at least one dataset (HipSci iPSC or ESC/iPSC additionally sequenced), much greater than expected (*p* = 1.15×10^−80^, binominal test). For rtQTLs with *p* ≤ 5×10^−8^, 85.6% (89/104) were validated (*p* = 3.75×10^−74^). Among the iPSC rtQTLs tested, 31.7% (303/955) were validated in ESC (*p* << 2.2×10^−16^). For iPSC rtQTLs with *p* ≤ 5×10^−8^, 82.3% (149/181) were validated (*p* << 2.2×10^−16^).

### A promoter-enhancer logic of replication timing regulation

We first used rtQTLs to examine the *cis*-regulatory logic of human replication timing. We observed that a subset of rtQTLs are distal to their associated genomic region (Fig. 1M), and that in many regions, separate rtQTLs clustered in close proximity. This suggests that multiple DNA sequences, local and distal, could interact to affect the replication timing of a given locus. We identified 318 cases, encompassing 803 rtQTLs, where at least two, and strikingly, up to seven rtQTLs were associated with the same region, each providing additional explanatory power (Fig. 2, A–D). We call these “multi-rtQTL” regions and refer to the strongest rtQTL as the “primary”, while all other rtQTLs are “secondary”. In some cases, one rtQTL quantitatively influenced replication timing, while several rtQTLs together explained the actual presence of active initiation (Fig. 2C). Thus, replication initiation is regulated along a continuum, one extreme of which is no activity at all despite the presence of a potential initiation site.

**Figure 2.**
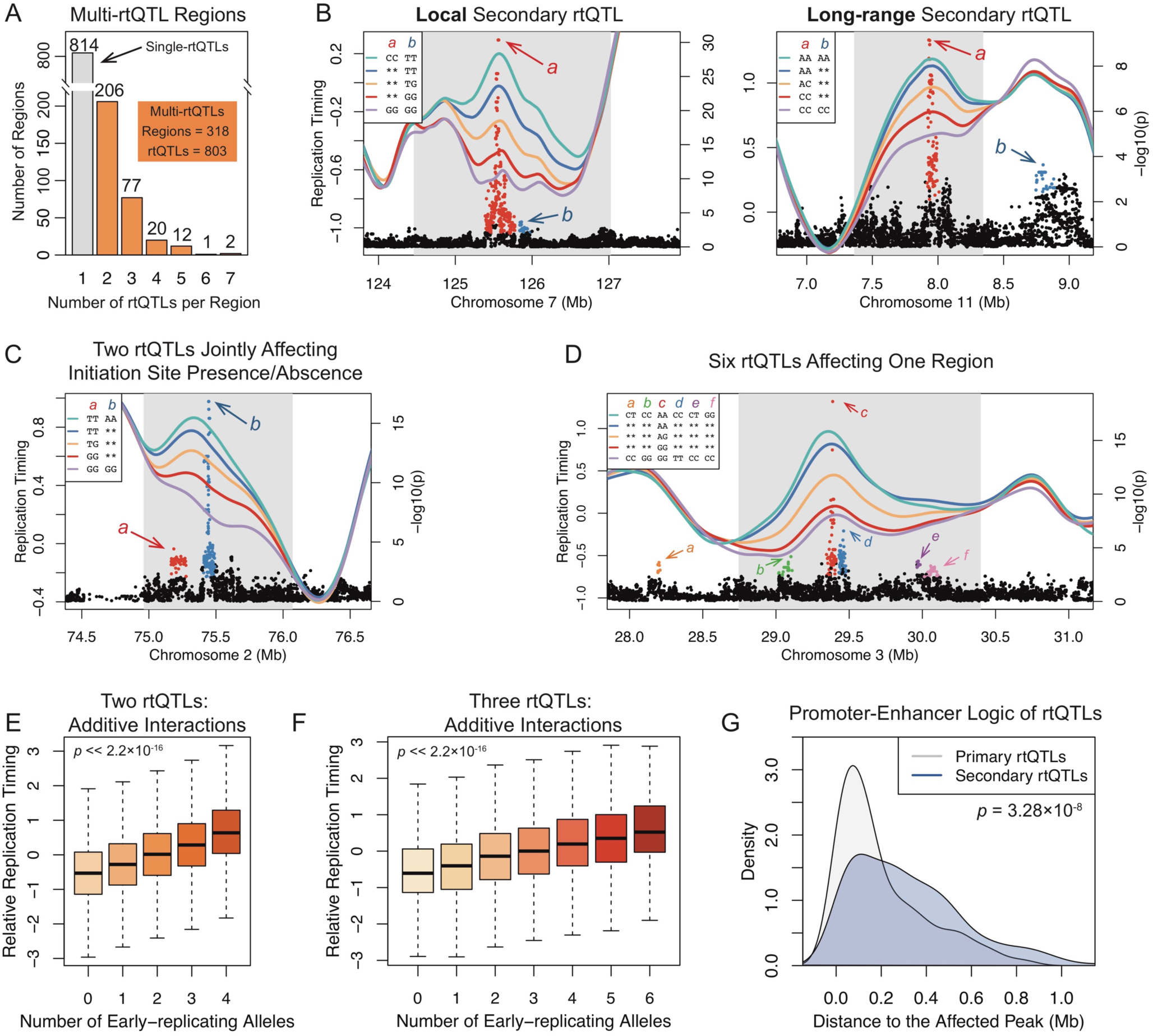
Multiple DNA Sequences Interact to Regulate Replication Timing. (A) Hundreds of regions are controlled by multi-rtQTLs. (B, C) Two rtQTLs affecting the same region. Blue, yellow, and red lines represent one rtQTL. Purple and green lines represent the mean replication timing of individuals carrying the late- or early-replicating genotypes, respectively, at both rtQTLs. Considering both rtQTLs explains a larger fraction of variation (green lines are higher than blue lines; conversely for purple/red lines). Asterisks (in legends): any genotype at this rtQTL. In panel C, the GG/GG combination of alleles is associated with complete loss of initiation activity. (D) A replication initiation site associated with six rtQTLs. Each rtQTL was significant even after conditioning on all other five rtQTLs in the region. (E, F) rtQTLs exert additive effects. All regions with two (E) or three (F) rtQTLs were pooled; replication timing is linearly correlated to the number of early-replicating alleles. (G) Multi-rtQTLs conform to a “promoter-enhancer” logic, primary rtQTLs being closer to the affected replication timing peak than secondary rtQTLs.

We directly tested for interactions between primary and secondary rtQTLs at regions that harbored two rtQTLs, hence between zero and four early-replicating alleles. We further pooled all genomic regions containing three or four rtQTLs and evaluated the relationship between the number of early-replicating alleles and replication timing of the associated regions. Replication timing showed a linear relationship with the number of early-replicating alleles (linear regression *p* << 2.2×10^−16^; Fig. 2, E and F), and none of the individual regions showed evidence for synergistic interactions between rtQTLs. This suggests that primary and secondary rtQTLs additively affect local replication timing.

Of the 318 multi-rtQTL regions, 176 were associated with replication timing peaks. In 115 of these cases (65.3%), primary rtQTLs were closer to the peak than secondary rtQTLs (Fig. 2G, *p* = 3.28×10^−8^). This resembles eQTLs (expression QTLs), in which primary eQTLs show stronger enrichment at promoters, while weaker eQTLs are enriched at enhancers^24^. Also in resemblance to enhancers and promoters, primary and secondary rtQTLs tended to cluster in nuclear space (based on Hi-C data) more than expected by chance (*p* = 9.73×10^−3^, *Z*-test). Drawing from this analogy, we propose that rtQTLs may follow a logic akin to promoters and enhancers, in which primary rtQTLs function as main *cis*-acting regulators of replication initiation, while other sequences, marked by secondary rtQTLs, serve as distal regulatory elements that fine-tune the replication dynamics of a given region.

### A histone code for DNA replication initiation

We next utilized the basepair-resolution sequence-specificity of rtQTLs to investigate the molecular determinants of DNA replication timing. We initially considered rtQTL locations *per se*, independently of allelic variation. Since extensive epigenetic data was available for seven of the hESC lines in our dataset, we focused this analysis on hESCs and used iPSCs for validation. Consistent with previously described correlations between early replication and open chromatin^3^, rtQTLs were enriched for active chromHMM states including enhancers and transcription start sites (although they were not specifically associated with genes; Fig. S2), DNase I hypersensitivity sites (*p* = 2.62×10^−8^, 4.11×10^−19^ in iPSCs), and H2A.Z sites^25^ (*p* = 6.69×10^−4^; *p* = 3.20×10^−17^ in iPSCs). rtQTLs also significantly overlapped with 24 histone marks (25 in iPSCs), of which 20 were active marks (Fig. S2). The majority of these histone marks were acetylations, including several not linked to replication timing before, for example, H2BK120ac, H2BK12ac and H2BK20ac. H3T11ph was also consistently enriched at hESC and iPSC rtQTL sites, as so were, modestly, methylated forms of H3K4.

Of note, the histone mark enrichments were modest, and each present at between 165 to 541 (median: 429) of 608 hESC rtQTLs (median: 542 of 1,167 iPSC rtQTLs), while each rtQTL overlapped 20 histone marks on average. We surmised that this abundance of histone modifications may be suggestive of combinatorial regulation. To test this, we systematically searched for combinations of histone marks with stronger enrichments at rtQTLs when considered jointly (Methods). We identified 152 combinations of two overlapping histone marks that were more enriched than the individual marks. We further identified 128 co-enriched three-mark combinations, 72 four-mark combinations, and 13 five-mark combinations (enrichment *p*-values: 2.42×10^−37^–1.09×10^−45^), at which point no further improvements in enrichment were obtained (Fig. 3A). Importantly, since these enrichments controlled for replication timing, they were not identified because they mark early-replicating regions, but because they specifically mark rtQTL locations, and, by inference, replication initiation sites.

**Figure 3.**
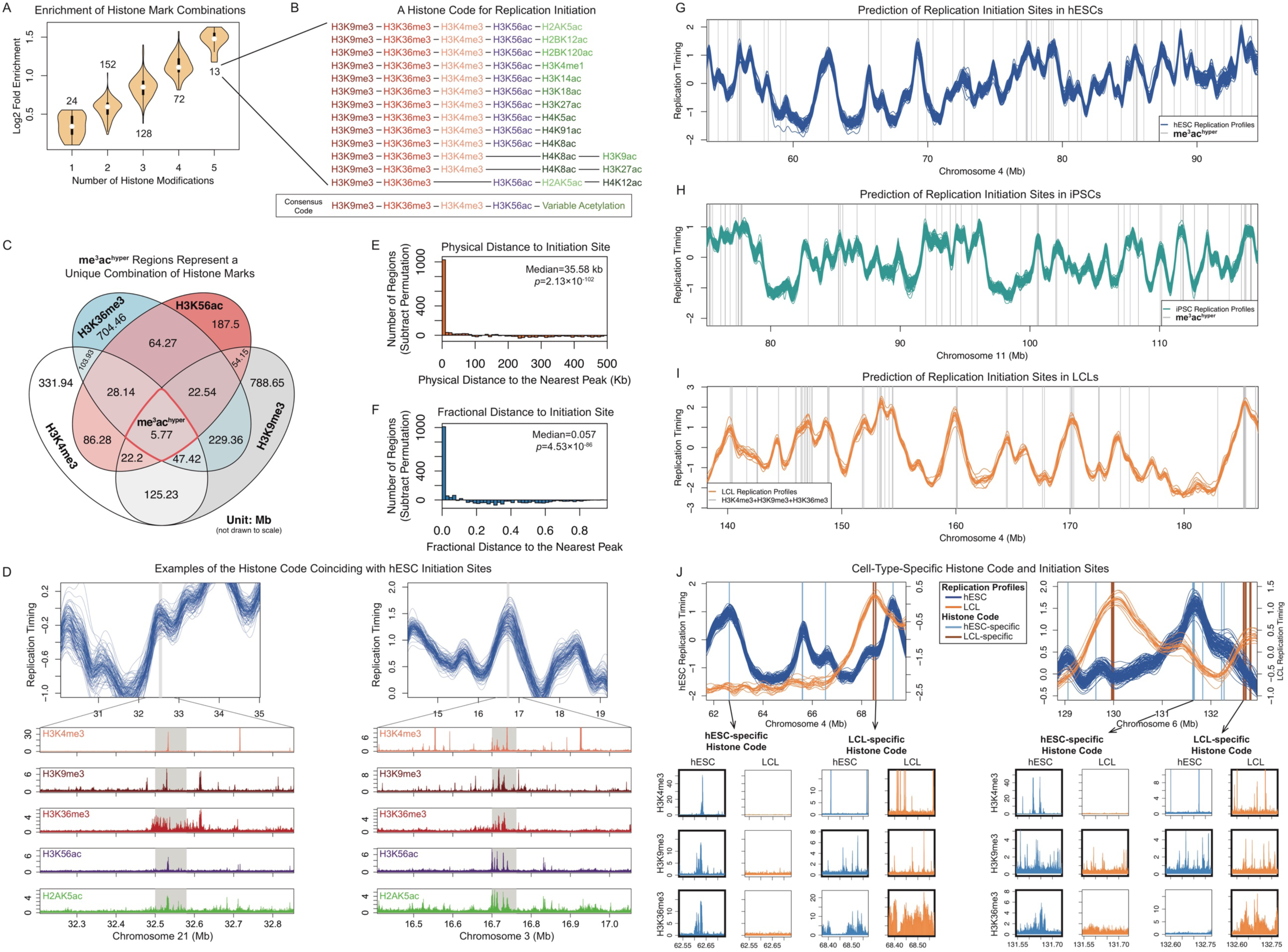
A Histone Code for Replication Initiation. (A) Iterative identification of histone mark combinations enriched at rtQTLs. Shown are enrichment distributions; the number of combinations in each category is indicated. Fold-enrichment increases gradually and is maximal for five-mark combinations. (B) A histone code for human replication initiation. The 13 combinations of five histone marks converged to a consensus “code”. (C) The histone code represents a rare combination of both active and repressive histone marks. me^3^ac^hyper^ regions comprised 0.7–3% of the regions that carry the individual histone marks. (D) Examples of histone mark combinations (Roadmap Epigenomics imputation)^39,40^ coinciding with replication timing peaks not identified as rtQTLs. (E, F) Distribution (after subtraction of permutations) of physical (E) and fractional distances (F) of the me^3^ac^hyper^ locations to the nearest replication timing peak. (G) Combination of histone marks (gray, me^3^ac^hyper^ locations) predict replication initiation sites in hESCs. (H, I) Histone code locations (gray vertical lines) correspond to replication timing peaks in iPSCs (H) and LCLs (I). (J) Cell-type-specific histone code locations mark cell-type-specific replication initiation sites. At regions with distinct replication timing profiles for hESCs and LCLs, LCL (hESC)-specific replication timing peaks are predicted by LCL (hESC)-specific histone code locations. Lower panels: initiation sites coincide (thick borders) with all three histone trimethylation marks in the cell type in which they are active, but with one or none of the marks in the cell type in which they are inactive.

Strikingly, all 13 combinations of five histone marks contained the trimethylation marks H3K9me3 and H3K36me3, and 12 of the combinations also contained H3K4me3. In addition, all 13 combinations included at least one histone acetylation mark. H3K56ac was included in 11 of the combinations, while the additional acetylations occurred on variable histone residues (Fig. 3B). Further analysis indicated that various acetylation marks often coincided with the five histone mark combinations, e.g., in 70.8% of the cases, 11 or more acetylation marks co-occurred at the location of a five-mark combination. We term this combination of <u>three H3 tri</u>metylations together with hyperacetylation the “me^3^ac^hyper^ histone code”. Genome-wide, there were 6,670 such locations in hESCs. They covered a median of 635 bp and cumulatively encompassed 0.24% of the genome, thus they represent specific, localized genomic sites.

Importantly, when considered individually, the implicated histone modifications only showed weak enrichments (Fig. S2C). H3K9me3 and H3K36me3, in particular, showed marginal or no enrichment at rtQTLs. H3K9me3 is a marker of heterochromatin (although has been observed in the bodies of actively transcribed genes)^26^, while H3K4me3 marks gene promoters and H3K36me3 is typically present in gene bodies. These histone trimethylations are largely mutually exclusive. However, in rare cases, they coincide in the same genomic locations (Fig. 3, C and D) to form a previously undescribed bivalent chromatin that is not specifically associated with genes or gene expression (e.g., only 5.2% of me^3^ac^hyper^ regions overlapped the TSS of an active gene). It is in these rare locations that rtQTLs tend to be present.

The identified histone mark combinations have been previously linked to the recruitment of components of the replication machinery to DNA. Histone H3 trimethylations on lysines 4, 9 and 36 have been shown to exert a cross-talk that serves as an “epigenetic addressing system” for site-specific replication initiation^27,28^. They recruit KDM4 and KDM5 family histone demethylases that directly interact with, and/or are required for recruitment to DNA of MCM, PCNA, DNA polymerases and other replication factors^27–31^. H3K4me3 also synergizes with flanking H3K9ac and H3K14ac (both identified as part of the me^3^ac^hyper^ histone code) to recruit chromatin readers to DNA^32^. Another study showed that histone hyperacetylation synergizes with H3K9me3 to promote early replication of otherwise late-replicating mouse chromocenters^33^. In turn, acetylated histones have been shown to recruit replication initiation factors including TICRR/TRESLIN, ORC and MCM, via mediators such as BRD2, BRD4 and the histone acetyltransferase HBO1 (<u>h</u>istone acetyltransferase <u>b</u>inding to <u>O</u>RC)^14,34–36^. In particular, HBO1 promotes MCM loading by acetylating H4 on lysines 5, 8 and 12, and subsequently promotes origin activation by acetylating H3K14^37^; we identified all of these acetylations as part of the me^3^ac^hyper^ combinations. Moreover, H4K12ac, the most strongly enriched mark at rtQTLs, is a preferred target of HBO1 at replication origins^34,36^. These biochemical evidence provide a plausible explanation for the combination of histone marks being associated with replication initiation activity.

Taken together, we identified a combination of histone marks, consisting of three trimethylated H3 residues (H3K4me3, H3K9me3, H3K36me3) together with H3K56ac and broadly hyperacetylated chromatin that consistently coincide with rtQTLs. To further test the involvement of this histone “code” in replication initiation, we analyze below its association with: (1) replication timing peaks in general (independent of rtQTLs); (2) replication timing peaks in other cell types; (3) replication timing peaks that vary between cell types; and (4) replication timing variation among individuals at rtQTLs.

**Figure S2.**
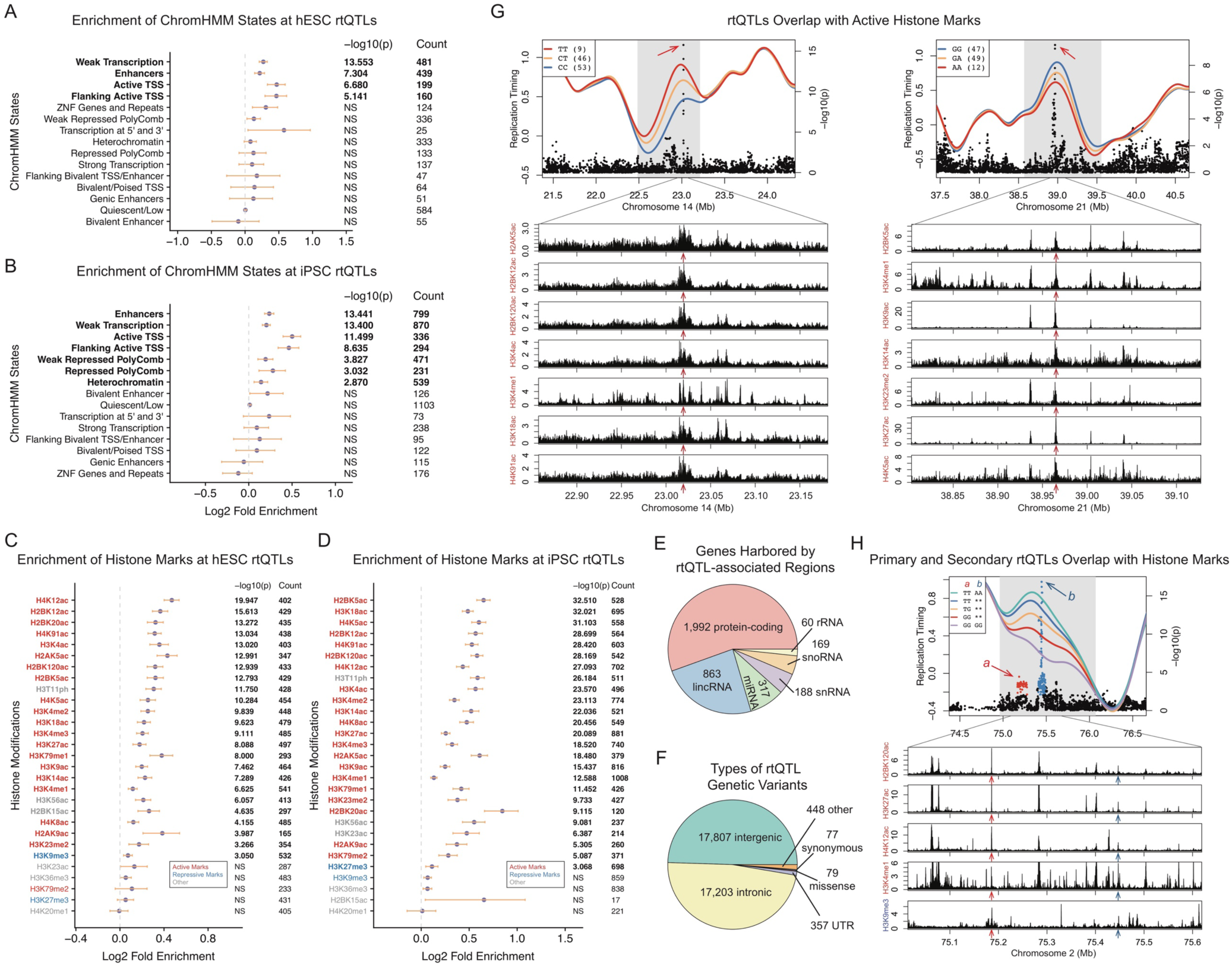
rtQTLs are Enriched for Active Chromatin States and Histone Marks. (A, B) Enrichment of chromHMM chromatin states at rtQTLs identified in hESCs (A) or iPSCs (B). Orange bars: 95% confidence intervals. NS: not significant at Bonferroni-corrected *p* = 0.05. (C, D) Enrichment of histone marks at hESC (C) and iPSC (D) rtQTLs. Similar to panels A and B. (E) Breakdown of gene types located within rtQTL-associated regions. The number of genes in rtQTL-associated regions was significantly lower than expected (*p* = 4.85×10^−17^, *Z*-test) and these genes were not enriched for any gene ontology term^38^. (F) Breakdown of functional annotations of rtQTL genetic variants. (G) rtQTLs colocalize with active histone modifications. The bottom panels show ChIP-seq tracks of various active histone modifications in hESC. Imputed histone tracks^39,40^ from the Roadmap Epigenomics Project were used for plotting. Red arrows: locations of the rtQTL variants indicated in the top panels. (H) A multi-rtQTL region (same as Fig. 2C) at which both the primary and secondary rtQTLs overlap with various active histone marks.

### The histone code predicts replication initiation sites across cell types

We considered whether a histone code could be a general property of replication initiation sites, revealed by leveraging the base-pair resolution of rtQTLs, but not limited to rtQTLs. We therefore tested whether the histone code also associated with the larger number of replication timing peaks (found in > 10% of the samples) not identified as rtQTLs (81.5% of all peaks). While the probabilities of having a peak near the 24 individually-enriched histone marks were significantly greater than expected (one-tailed Wilcoxon rank-sum *p* = 4.96×10^−17^) and were greatest at the actual histone mark sites (Fig. S3, A and B), individual histone marks are very common in the genome and insufficient for predicting peaks. Combining histone marks gradually increased their association with peaks, up to the five-mark combinations, which were significantly more likely than expected to coincide with peaks (*p* = 4.10×10^−10^). me^3^ac^hyper^ sites had an even higher likelihood of overlapping peaks (Fig. S3, A–C). The distances of me^3^ac^hyper^ regions to the nearest peak were significantly shorter than permutations (Fig. 3, E and F). Of all me^3^ac^hyper^ sites, 57.3% corresponded to replication timing peaks within 100 kb (positive predictive value; *Z*-test *p* << 2.2×10^−16^; the median inter-peak distance was 971.2 kb); 41.7% were less than 10 kb from a peak. Conversely, 70.8% of peaks were located within 100 kb of predicted regions (sensitivity; *p* = 1.03×10^−93^); 60.3% were less than 10 kb from predicted regions. We further evaluated prediction performance of the me^3^ac^hyper^ regions visually (Fig. 3G) and with ROC curves (Fig. S3D). Peaks predicted by histone marks replicated earlier than other peaks (median: 0.61 vs. 0.14, Wilcoxon rank-sum *p* = 6.58×10^−53^) and were locally more prominent (timing difference compared to flanking valleys, median: 0.32 vs. 0.18, *p* = 1.19×10^− 17^). Consistently, the replication profiles surrounding me^3^ac^hyper^ sites formed a sharp peak (Fig. S3E). The histone code was substantially more specific and matched replication timing profiles much better than DNase I hypersensitivity (Fig. S3G), which was previously suggested to explain 87% of replication timing profiles^41^. Taken together, the combinations of histone marks that are enriched at rtQTLs predict ∼70% of initiation site locations across the genome, even for those sites without rtQTLs, and particularly for the early and most prominent initiation sites. These histone mark combinations may thus promote replication initiation not just at specific genomic loci, as previously proposed^27,28,42^, but across a large fraction of the genome. We note, though, that some replication timing peaks did not co-localize with histone code locations, thus there must be additional mechanisms independently specifying replication initiation sites, underscoring the complexity of mammalian replication initiation.

An even more rigorous test of the five-mark combinations being indicators of replication initiation is whether they could predict the location of replication timing peaks in other cell types. Examining both iPSCs and lymphoblastoid cell lines (LCLs)^43–45^, we found that the histone code can predict initiation sites as accurately and specifically as in hESC (Fig. 3, H and I), and similarly associates with early replication (Fig. S3E). In particular, LCLs have epigenetic and replication timing landscapes that are distinct from those of hESC (and iPSCs). In genomic regions at which LCL and hESC replication timing differed, LCL-specific histone code locations corresponded to LCL-specific initiation sites, and *vice versa* for hESCs (Fig. 3J). Predicted cell-type-specific initiation sites resided in early-replicating genomic regions in the corresponding cell type, but not in other cell types (Fig. S3F). Thus, the histone code characterizes and predicts cell-type-specific replication initiation.

**Figure S3.**
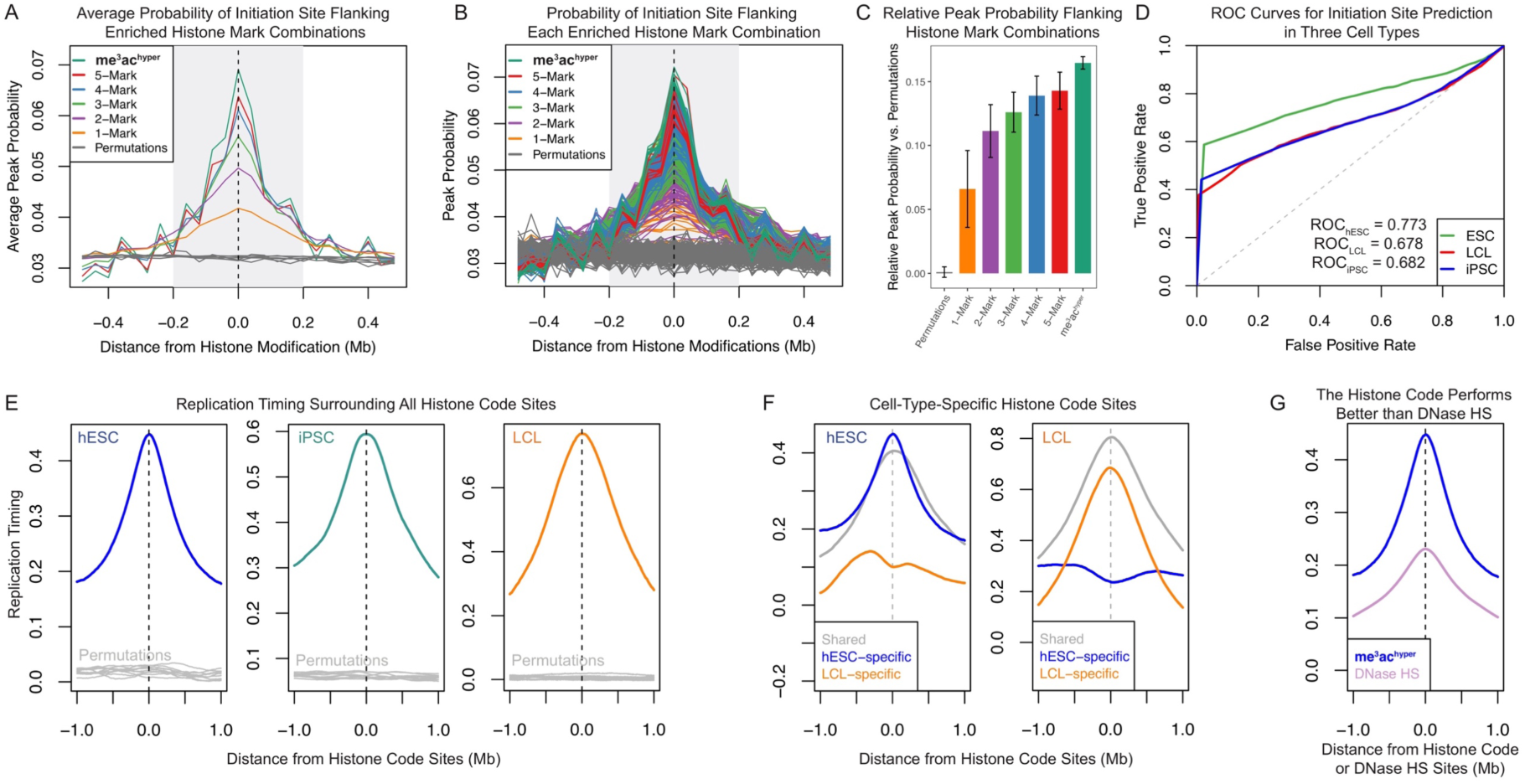
Further Support for a Histone Code for Human DNA Replication Initiation Sites. (A) Histone mark combinations correspond to replication initiation sites. The probability of having an initiation site increases with proximity to the histone mark combinations (gray shade), peaks at the actual histone mark sites, and scales with the number of marks. (B) Probability of having an initiation site as a function of distance from histone marks (in 40 kb bins), similar to panel A but for each individual histone mark combination (as opposed to the averages of all combinations of a given number of marks). (C) Normalized cumulative probability of initiation sites being present within 200 kb (i.e., area under the curve, gray shade in panel A) of individual histone marks or combinations thereof. The probabilities were normalized based on permutations by subtracting the permutation mean. Replication initiation sites are increasingly enriched as the number of histone marks increases. Error bar: standard deviation. Error bars: standard deviation. (D) ROC curves show the strength of the histone code for predicting replication initiation sites in various cell types. Diagonal lines represent random guesses. For all three ROC curves, the area under the ROC curve (AUC_ROC_) is significantly larger than random permutations (all *Z*-test *p* << 2.2×10^−16^). (E) Cumulative replication timing profiles surrounding histone code locations suggest that they coincide with locally early replication across cell types. For LCLs, only methylation marks were available. Gray lines: ten permutations. (F) Cumulative replication timing profiles in hESCs and LCLs surrounding histone code locations found in both cell types (gray), LCLs only (orange), or hESCs only (blue). Histone code locations predict replication initiation patterns in a cell-type-specific manner. (G) The histone code performs better at predicting replication timing peaks than DNase hypersensitivity (HS) sites. Cumulative replication profile centered at histone code locations (blue) is sharper and higher than that centered at DNase HS sites (purple). In addition, there are > 99,000 DNase HS sites in the genome, totaling > 304 Mb of sequence (i.e., ∼10% of the genome; in contrast to the histone code covering 0.24% of the genome), which provides very low positive predictive value and resolution for predicting individual replication initiation sites.

### Co-variation of replication timing and histone modifications reveals combinatorial control of replication timing

The previous analyses considered rtQTL locations *per se*. However, since rtQTLs represent replication timing variation among individuals, their allelic differences provide a powerful opportunity to investigate molecular mechanisms controlling replication timing. In particular, given that specific histone marks associate with replication initiation, we predicted that rtQTL SNP alleles will be associated with variation in the abundance of these marks among individuals.

We took an unbiased approach using seven hESC lines with both replication timing and histone modification data (Methods). Cell lines carrying early-replicating genotypes at rtQTLs were more likely than individuals with late-replicating genotypes to harbor active histone marks and chromHMM states at those rtQTL sites (Fig. 4 and S4). Across individuals and genomic sites, eight histone modifications were consistently present in individuals with rtQTL alleles indicative of early replication. Of those, seven were acetylations, consistent with histone acetylation promoting early replication^3,13–17,34,36^. Of the 12 acetylation marks that are part of the replication initiation histone code, nine individually associated with early-replicating rtQTL genotypes (five of which reached statistical significance). We also identified seven modifications that consistently coincided with late replicating alleles, of which six were methylation marks (Fig. 4A); Thus, histone methylation emerges as being generally repressive for replication.

**Figure 4.**
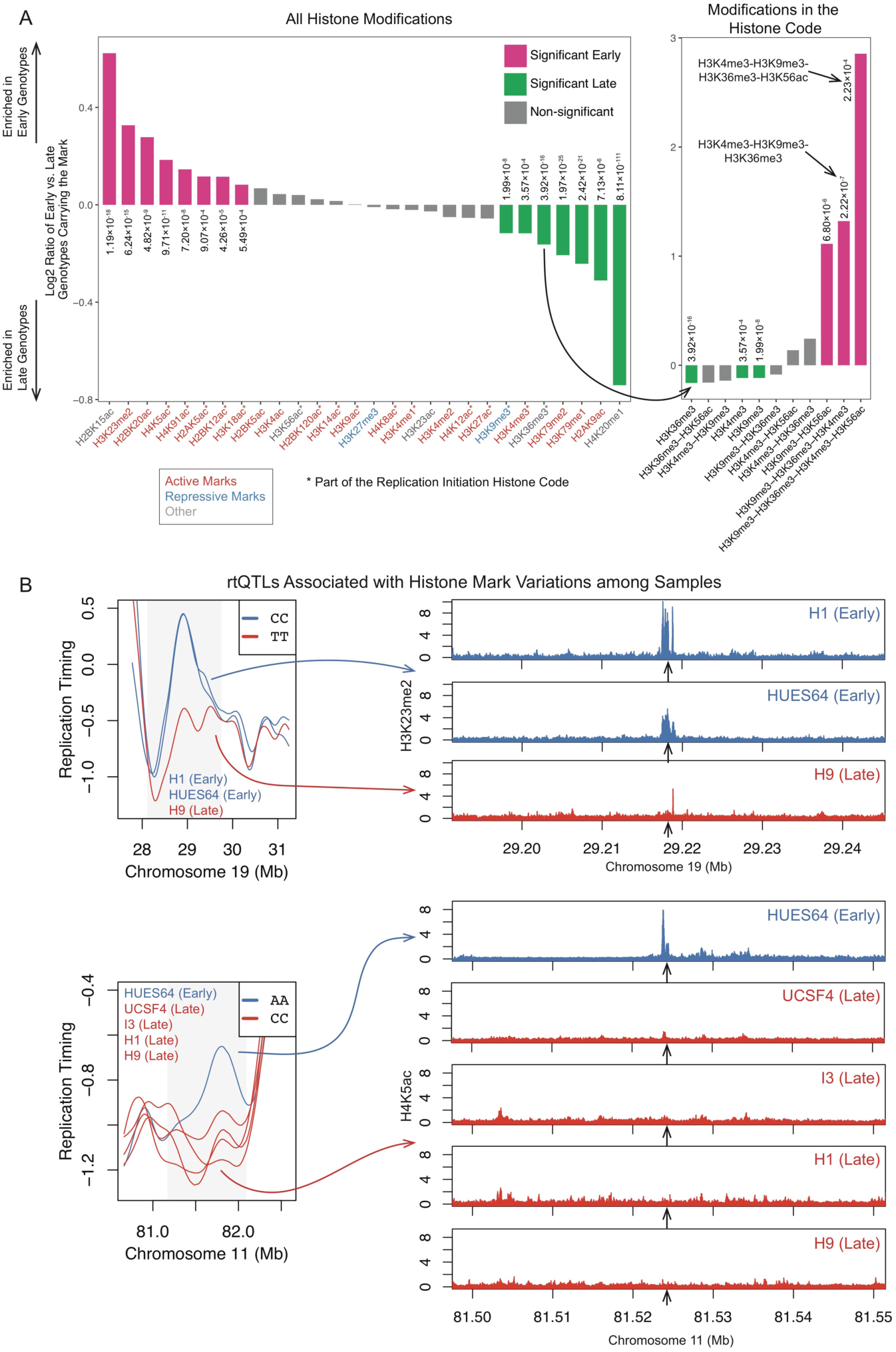
Histone Marks Affect DNA Replication Timing. (A) Association of rtQTL genotypes with individual (left panel) or combinations (right panel) of histone marks. Positive (negative) values indicate that individuals with early (late)-replicating genotypes are more likely to carry a histone mark at those rtQTL sites. Right panel: while individual H3 methylation marks associate with late replication, the H3K4me3-H3K9me3-H3K36me3 combination is strongly associated with early replication, and even more so when combined with H3K56ac. Note the different Y scale. (B) Examples of rtQTLs associated with histone mark variations. Replication timing and corresponding histone ChIP-seq tracks for individual cell lines homozygous for the early- or late-replicating alleles. Early replication correlates with the presence of the specified histone marks.

**Figure 5.**
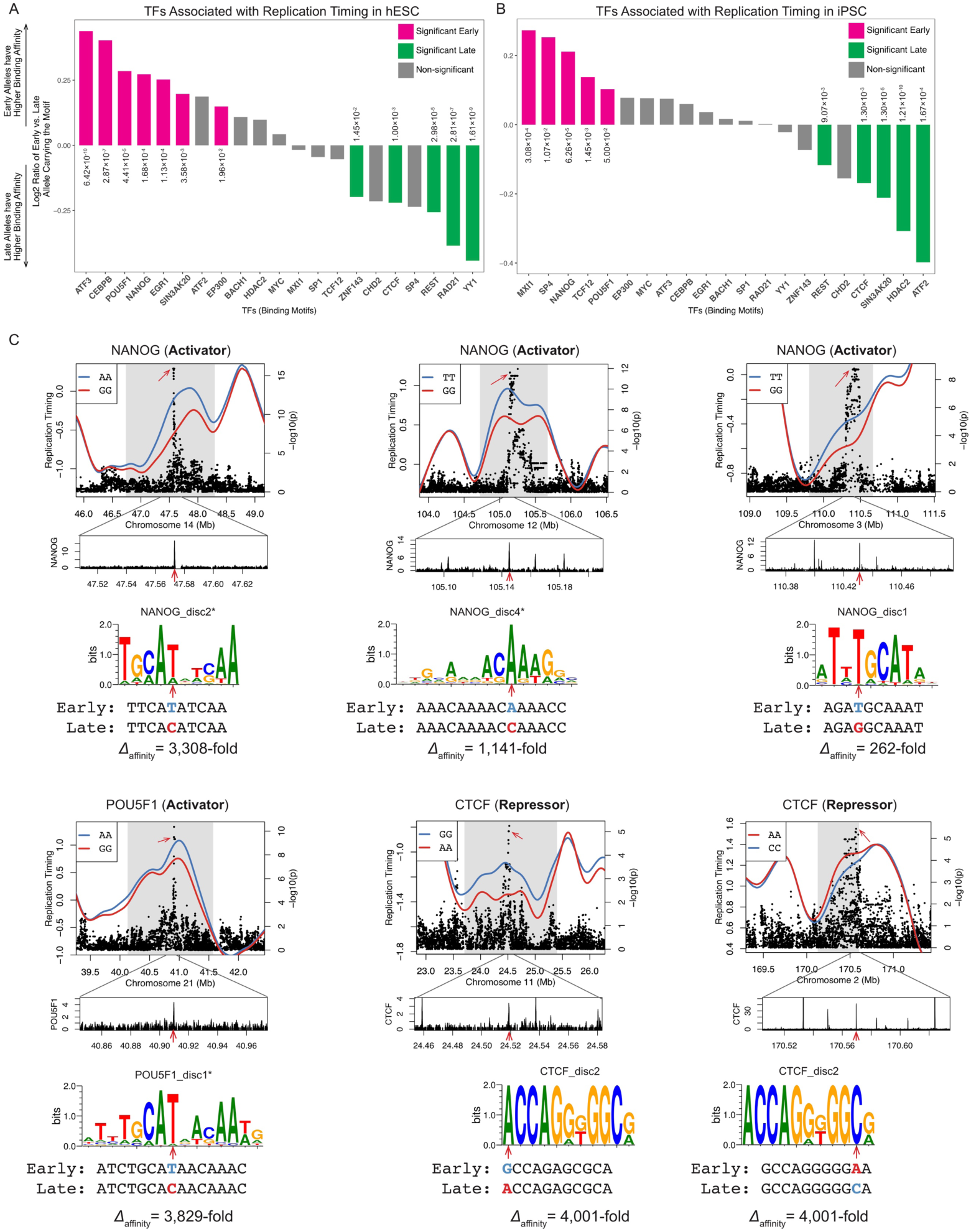
rtQTLs Affect Replication Timing by Altering TF Binding Motifs. (A, B) Binding of TFs such as OCT4 and NANOG promotes earlier replication, while binding of CTCF, REST and other factors is associated with later replication in hESCs (A) and iPSCs (B). Chi-squared test, FDR <10%. (C) Examples of rtQTLs altering binding affinity of TFs that function as replication activators or repressors. Heterozygous profiles are not plotted. Center panels: ChIP-seq tracks. Lower panels: sequence logos of the motifs containing the rtQTL SNPs, motif names, and changes in binding affinity (calculated based on motif scores). Asterisk indicates that the motif was on the negative strand and the sequence shown is the reverse complement. Red arrows: locations of the rtQTL SNPs. For activators, the rtQTL allele associated with early replication encodes an intact binding motif, while the allele associated with late replication abolishes the motif. Repressors have the opposite pattern: the early allele abolishes the motif.

Counter-intuitively, the histone code trimethylation marks (H3K4me3, H3K9me3 and H3K36me3) were individually more likely to be associated with late-replicating genotypes (Fig. 4A). In contrast, the combination of all three trimethylation marks was 2.5-times more likely to be carried by early-replicating than by late-replicating genotypes. Furthermore, a combination that also included H3K56ac was 7.24-times more likely to be carried by early-replicating genotypes (Fig. 4A). Thus, these marks appear to individually act as weak repressors of replication but act synergistically, in non-canonical ways, to strongly promote early replication. Taken together, the involvement of me^3^ac^hyper^ in replication initiation is supported by several lines of evidence: enrichment at rtQTLs (Fig. 3A); correspondence with replication timing peaks in general, and across several cell types (Fig. 3, D–I, Fig. S3, A–E); co-variation with cell-type-specific replication initiation patterns (Fig. 3J and Fig. S3F); and correlation with inter-individual replication timing variation (Fig. 4).

**Figure S4.**
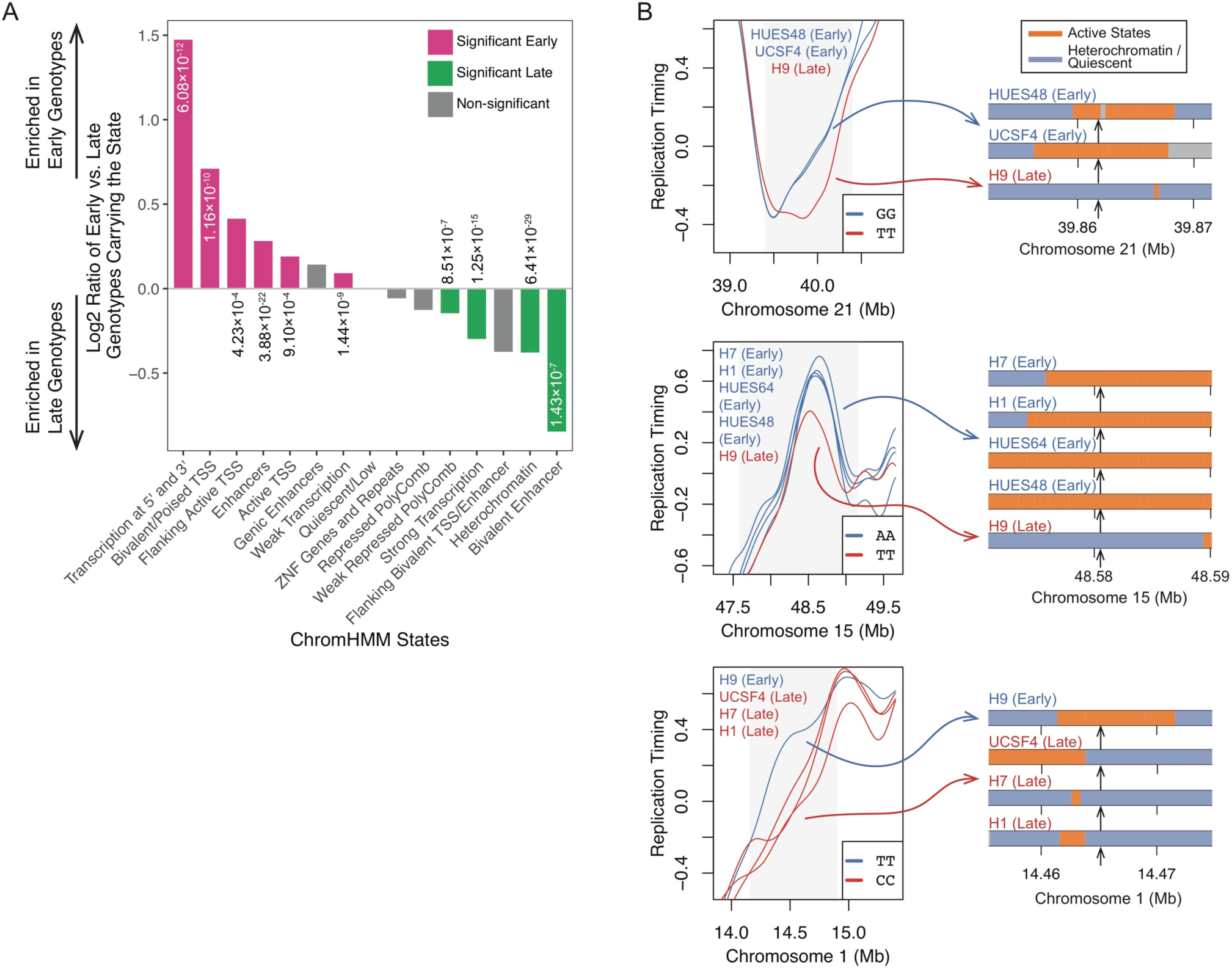
rtQTLs Impact Replication Timing by Affecting Chromatin States. (A) Associations of rtQTL genotypes with chromHMM states. Positive values indicate that the early-replicating genotypes are more likely to carry a given chromatin state, and *vice versa* for negative / late genotypes. (B) Examples of rtQTLs associated with chromatin states. The right panels show chromatin states flanking the rtQTL in the same cell lines. Orange: active states (TSS, enhancer, or weak transcription), blue: heterochromatin or quiescent states, gray: other states.

### DNA-binding factors modulate DNA replication timing

The above results indicate that *cis*-acting sequences, manifesting as rtQTLs, influence the positions and timing of replication initiation by associating with histone modifications. To identify additional factors that influence replication timing via *cis*-acting sequences, we analyzed the binding sites of 51 DNA binding factors in hESCs^43,46^. Binding of eight factors was significantly enriched at rtQTLs, including the main pluripotency factors SOX2, POU5F1 (OCT4) and NANOG, the latter two reproducible with available data in iPSCs (Fig. S5). Three chromatin remodelers, EP300 (P300), SP1, and RBBP5, were also enriched at rtQTLs. EP300 is a histone acetyltransferase that catalyzes at least six acetylation marks in the replication initiation histone code, including H3K56ac^47^.

Transcription factors (TFs) bind DNA in a sequence-specific manner at characteristic motifs. This offers an opportunity to test, at base pair resolution, whether TF binding affects replication timing at rtQTLs (Methods). Strikingly, OCT4 and NANOG had significantly higher binding affinity for early-compared to late-replicating alleles in both hESCs and iPSCs, while EP300 and ATF3 (Activating Transcription Factor 3, which is enriched at EP300 sites^48^), were linked to early replication at least in hESCs (Fig. 6, A and B). These associations appeared to be independent from gene expression, as they were reproduced for rtQTLs > 250 kb away from expressed genes. For these early-replication-associated TFs, the rtQTLs fell within the TF binding motifs such that a single base-pair change disrupted or even abolished binding; this was associated with delayed replication of the rtQTL-affected initiation site (Fig. 6C). An unexpected finding was rtQTL alleles with the opposite effect, i.e., higher binding affinity associated with later-replication. We infer that in these cases protein binding suppresses replication initiation (Fig. 6). These included CTCF, an insulator of topologically associated domains (TADs); REST(NRSF), a repressor of transcription^49^; ZNF143, which associates with the CTCF-cohesin cluster^50^; and at least in hESCs also RAD21 (part of the cohesin complex) and YY1, which co-localize with CTCF at TAD boundaries^51–54^. These associations were yet stronger when considering only motifs with biochemically confirmed TF binding when data was available (Methods). Taken together, we conclude that some rtQTL alleles alter DNA binding protein motifs, abolish a DNA binding site or generate a new one, and consequently alter DNA replication timing through specific protein binding. This analysis uncovers several new factors that can thus regulate DNA replication timing. In addition, different factors influence subsets of replication initiation sites, further illuminating the complex combinatorial landscape that controls human DNA replication timing. Finally, these results demonstrate how a single base-pair alteration could affect the replication timing of an extended genomic region.

**Figure S5.**
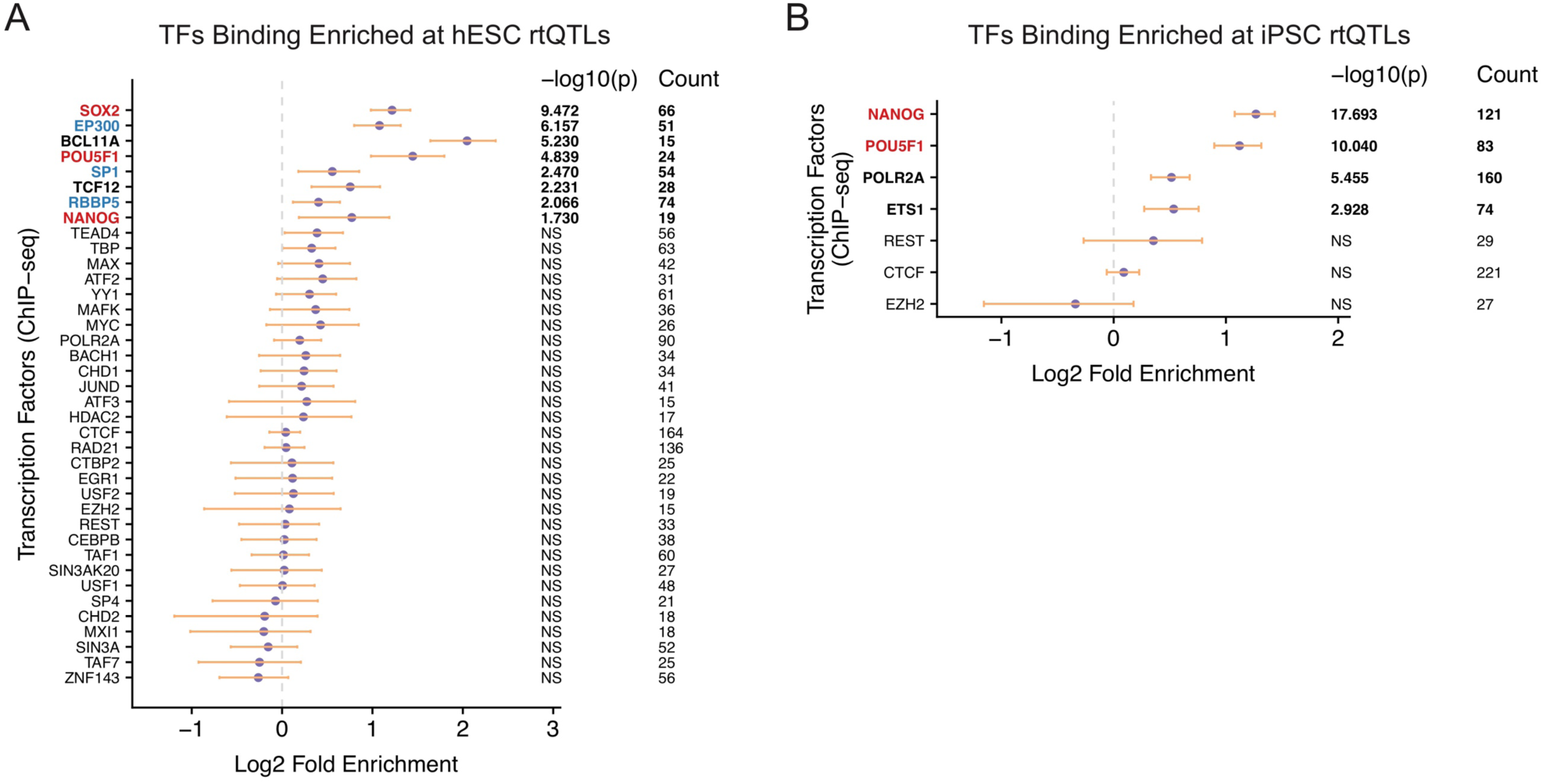
Enrichment of TFs at hESC (A) and iPSC (B) rtQTLs. rtQTLs are enriched at binding sites of central pluripotency factors (red) and chromatin remodelers (blue). NS: not significant at 10% FDR. Only TFs overlapping with at least 15 rtQTLs are plotted.

## Discussion

The spatiotemporal regulation of DNA replication, and its dependence on regulatory DNA sequences, are poorly understood. Here, we leveraged population-scale replication timing and genetic polymorphism data to perform the equivalent of millions of surgical genetic interrogations of replication timing determinants. This approach enabled us to identify an unprecedented number of precise sequence determinants of replication timing.

Studying chromatin structure at rtQTL sites revealed a histone code that accurately predicts replication initiation across cell types. This code represents non-canonical functions of histone H3 lysine methylations that form a previously undescribed bivalent chromatin state^55^ present at specific sites throughout the genome. Prior biochemical evidence supports an involvement of these histone marks in DNA replication initiation^14,27–32,34–37^. We propose that this histone code promotes local replication activity, although we do not necessarily imply that it marks the locations of replication origins *per se*.

rtQTLs further associated with inter-individual variation in histone marks and TF binding affinity. In many cases, several *cis*-acting sequences affected a region’s replication timing both proximally as well as distally. Altogether, we were able to assign at least one molecular determinant to 98.8% of rtQTLs, while two or more determinants were implicated in 93.9% of rtQTLs (Fig. S6). Replication timing determinants acted additively among nearby sequences, synergistically between histone modifications, and combinatorially across transcription factors. This system generates a continuum of replication activities: some epigenetic marks may contribute only modestly to replication activity, or even suppress it, yet can interact with other factors to ultimately promote robust early replication. Taken together, this study systematically reveals a complex, combinatorial landscape of genetic regulation of human DNA replication timing.

A recent study using CRISPR/Cas9-mediated deletions in mouse ESCs identified several interacting sequence elements responsible for early replication (“early replicating control elements”^12^). Consistent with our results, the identified elements bound P300 and pluripotency-related TFs. However, the specific features identified with deletions represented the properties of only 1.5% of rtQTLs. Instead, rtQTLs associated with replication throughout S phase (not just with early replication); some interacted with others while many did not; and there was no single DNA-binding factor that was always bound to them. rtQTL mapping reveals a much more complex picture of replication timing regulation than previous approaches were powered to uncover. Replication timing regulation emerges as a quantitative trait, requiring a quantitative genetics approach to elucidate its complex sequence underpinnings. rtQTL mapping in larger sample sets and additional cell types will further refine the details of replication timing regulation and reveal additional *cis*-acting sequences and their mode of action. In addition, rtQTL mapping refines the relationship between DNA replication timing and gene expression and reveals influences of replication timing on personalized mutational landscapes and on human phenotypes including disease risk^4,22^ (our unpublished results).

Our findings draw corollaries between replication timing regulation and classical concepts of gene expression regulation: promoter/enhancer logic, activators and repressors, and a histone code. Thus, replication and transcription regulation appear to be based at least in part on similar principles and building blocks. Replication timing is robustly encoded in DNA, yet multiple DNA sequences dictate DNA replication combinatorially via chromatin effectors. The replication timing program of the human genome emerges as being sequence-dependent, without being sequence-specific.

**Figure S6.**
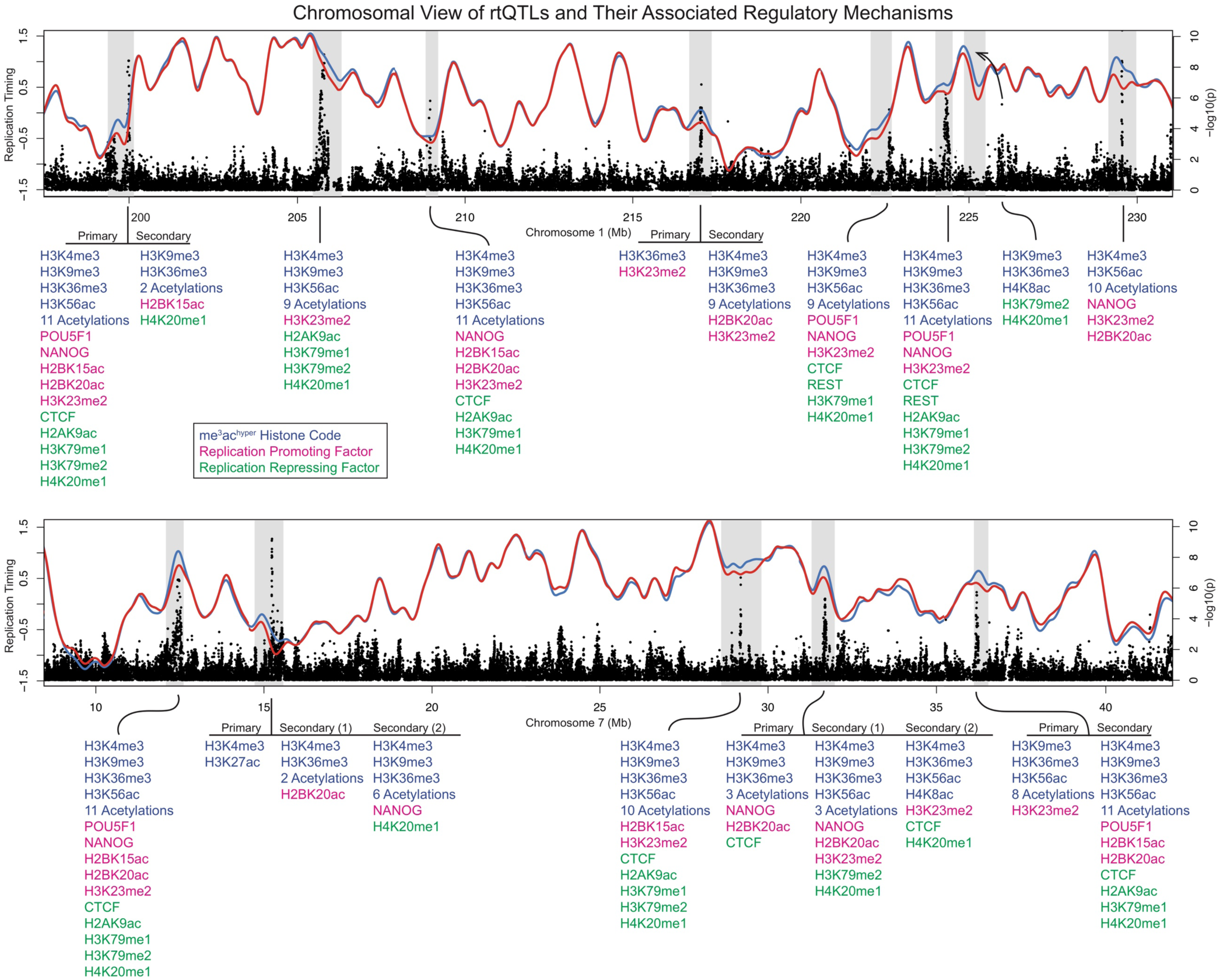
rtQTLs Regulate Replication Timing via Numerous Activating and Repressing Effectors. Different combinations of TFs and histone marks exert positive and negative effects on subsets of replication initiation sites. Both examples show 10 ESC rtQTLs spanning a ∼30-Mb region (on chromosomes 1 and 7). The blue and red lines are mean replication profiles of individuals carrying the early- and late-replicating genotypes, respectively. The rtQTL at 225 Mb of chromosome 1 exerts a long-range effect (arrow). Histone marks and TFs overlapping rtQTL genetic variants are shown below. They include positive (magenta) and negative (green) determinants of replication timing (Fig. 4 and 5), and instances of the replication initiation histone code (blue, Fig. 3).

## Data Availability Statement

Data of hESC and iPSC lines sequenced in this study were desposited in dbGaP (accession number: phs001957).

## Code Availability Statement

Computer codes used in this study are available from the corresponding author upon request.

## Supplementary Materials

### Materials and Methods

#### Whole-genome sequence data

Whole-genome sequence data and genotype calls for 121 hESC lines were obtained from Merkle et al. (submitted). We denote these as “Merkle_batch1”. Nine additional hESCs were sequenced in another batch from Merkle et al. (submitted), denoted as “Merkle_batch2”. We further used whole-genome sequence data from 326 iPSC lines from the HipSci Project^19^ (ENA accession number: PRJEB15299), denoted as “HipSci”. We sequenced an additional 15 hESCs and 17 iPSCs (dbGaP accession number: phs001957), denoted as “in_house_hESC” and “in_house_iPSC”, respectively.

For the in-house datasets, DNA was extracted using the MasterPure Complete DNA and RNA Purification Kit (Lucigen). Sequencing libraries were prepared using the Illumina TruSeq PCR-free kit and sequenced on an Illumina HiSeq X Ten to ∼16-fold coverage with 150×2 paired-end reads. Sequencing was performed at GeneWiz (South Plainfield, NJ). No approval was needed for sequencing. Reads were aligned to GRCh37 using BWA, and genetic variants were called following the GATK Best Practices. Variants were filtered using GATK’s variant quality score recalibration, such that SNPs had a 99.9% sensitivity to true variants (HapMap and Omni 2.5M)^56^ and a 99.0% sensitivity to true indels (Mills / 1000 Genomes indels)^57,58^.

#### Inference of DNA replication timing

DNA replication timing was inferred by analyzing sequence read depth (corrected for GC content bias) in non-overlapping windows of 10 kb of uniquely alignable sequence using GenomeSTRiP ^4,59^. Among the 121 hESC lines from Merkle_batch1, five did not optimally thrive in culture, resulting in read depth profiles with low correlations to other samples; these cell lines were excluded from further analysis. We excluded 26 of the 326 iPSC lines from the HipSci dataset for the same reasons. As described below, further filtering were performed for Merkle_batch1 and HipSci datasets. Replication timing inference for the in-house datasets is described separately (see the “validation of rtQTLs” section below).

Replication timing windows falling under any of the following categories were filtered out in all cell lines: (1) spanning GRCh37 gaps; (2) overlapping structural variants (SV) with ≥ 1% MAF in the 1000 Genomes European individuals; (3) overlapping short CNVs (median size: 3.51 kb) identified directly in the analyzed cell lines (applicable for Merkle_batch1 only, Merkle et al., submitted); (4) having a median window copy number (across samples) at least 0.4 copies away from the median (across windows) of median copy number of all autosomal windows (across samples); and (5) the 25% / 75% percentile (across samples) copy number of the window was at least 0.4 copies away from the median (across windows) of 25% / 75% percentile copy number of all autosomal windows (across samples). Criteria 2 and 3 remove windows that are possibly affected by SVs or CNVs, while criteria 4 and 5 remove windows that had outlying copy number values across a significant portion of samples. Specific parameters for criteria 4 and 5 (as well as the filtering steps described below) were chosen after evaluation of data values that were consistent across chromosomal replication profiles and across samples. Altogether, 28,769 data windows (11.0% of all windows) were removed, leaving 232,027 windows after filtering for Merkle_batch1. For the HipSci dataset, 239,516 windows remained.

Replication timing windows falling under any of the following categories were filtered out in individual cell lines: (1) at least 0.6 copies away from the median (across windows) of median copy number of all windows (across samples); (2) at least 0.25 copies away from the median copy number of that replication timing window; (3) in a large CNV (median size: 3.02Mb) identified in that individual (applicable for Merkle_batch1 only); and (4) in a region suspected to be a large subclonal CNV (sub-integer change in copy number over a large region, usually an entire chromosome or a chromosome arm). These criteria were implemented to further remove outlier data points. Data after the above filtering steps is referred to as “filtered raw data”.

Processing of the X chromosome data was performed separately for males and females. For males, because they only carry one X chromosome, all the thresholds above were divided by two.

The filtered raw data was further normalized to *Z*-score (i.e., autosomal mean of zero and standard deviation of one) by subtracting the mean then dividing by the standard deviation of all data points, and smoothed using a penalized smoothing spline using the R `pspline` package with smoothing parameter 10^−16^. For each chromosome, we smoothed across gaps only if the gaps were shorter than 300 kb. Continuous genomic segments (between gaps) that were smaller than 300 kb were removed from further analysis. Data after the above normalization and smoothing is referred to as “smoothed data” (Fig. 1A) and was used in further analyses. The total length of replication timing windows in the smoothed data was 2,330.66 Mb for autosomes (referred to as the “analyzable genome”), 121.15 Mb for the X chromosome in females, and 121.19 Mb for the X chromosome in males. For analyses involving the analyzable genome, only autosomal rtQTLs were counted.

For correlation calculations involving sib pairs vs. non-sib pairs (Merkle_batch1) and cell lines derived from the same donor vs. different donors (HipSci), we used replication timing data from chromosomes 1 to 5. The Wilcoxon rank-sum test was used to assess significance. For the analysis regarding IBD segments in sib pairs, we first inferred pairwise IBD using TRUFFLE^60^, then binned the IBD segments into 2.5 Mb regions. The purpose was to minimize bias in correlation estimation because of variable IBD segment sizes. We calculated pairwise correlation in these regions, and assigned the estimate to one of three groups (IBD 0/1/2). ANOVA was used to assess significance of difference in average correlation among IBD 0/1/2 groups. For all box plots in this study, the center line represents median, box limits represent the first and third quartile, and the whiskers represent the maximum and minimum. Outliers as determined by the R `boxplot` function were not plotted.

#### Identification of replication timing peaks

We identified peaks in the Merkle_batch1 dataset. For each sample, peaks were identified in the replication timing profile as local maxima. Peaks across all samples were then clustered using agglomerative hierarchical clustering in MATLAB (functions `linkage` and `cluster`) with a distance threshold of 200 kb, which yields a list of peak clusters, each containing one or more peak locations. When a cluster contained multiple peaks from the same sample, the peak closest to the cluster center was retained and all other peaks from the sample were dropped. For each peak cluster, the boundary was defined as the full range of peak locations in this cluster. We only used peak clusters that contained peaks from more than 10% of the samples.

#### Identification of replication timing variants

We searched for replication timing variants using the Merkle_batch1 and HipSci datasets. We expect genomic regions with strong replication timing variation to have greater standard deviation (SD) across samples, compared to average genomic regions. We calculated SD across samples for each replication timing window across the genome. Since local maxima will indicate the highest regional SD values, we called peaks in SD across the genome. To prevent calling peaks at single outlier data points, we first smoothed the SD curve. Then, we removed peaks that were below a SD threshold equal to the mean of the genome-wide SD distribution. We performed pairwise *t*-tests on pairs of samples for replication timing difference on 500 kb windows centered at the remaining SD peaks. For example, in the Merkle_batch1 dataset, 2,154 windows were tested, of which 1,785 (82.7%) were significant at a Bonferroni-corrected significance threshold of *p* = 4×10^−9^. These significant windows were extended by testing adjacent 200 kb windows, sliding 100 kb at a time, until there were no longer any significantly different cell line pairs. After the extension step, SD peaks in close proximity occasionally resulted in overlapping replication timing variants. In these cases, if the correlation of replication timing across samples at the SD peaks was greater than 0.9, we merged these variants. Otherwise, adjacent variants were separated at the valley between SD peaks. Last, replication timing variants driven by less or equal than 1% of the samples were removed. This resulted in a total of 1,489 and 1,837 replication timing variants in the Merkle_batch1 and HipSci dataset, respectively.

#### Data processing prior to rtQTL mapping

##### Sample selection

We performed principal component analysis (PCA) on the genotypes of the hESC lines, using the 1000 Genomes Phase 3 European, East Asian, and African samples as references. Eight samples appearing to have non-European ancestry (admixed or East Asian) were removed from rtQTL mapping, leaving 108 individuals for further analysis. PCA was performed using the `SNPRelate` package in R ^61^. We also performed PCA with the HipSci dataset, and confirmed that all samples were of European ancestry. A total of 192 unrelated samples in the HipSci dataset were used for rtQTL mapping. While we kept sib pairs in the ESC dataset, all rtQTLs in ESC were reproducible (at nominal *p* < 0.05) when using only unrelated samples.

##### Genotype imputation

Imputation was performed with IMPUTE2 ^62^ using the 1000 Genomes Project Phase 3 reference panel and default parameters. Variants with minor allele frequency (MAF) ≤ 1% in Europeans or Americans were not used for imputation. Imputed variants with average genotype probability ≥ 80% were used in subsequent analyses.

Prior to rtQTL mapping, we filtered out variants that had MAF < 5%, were non-biallelic, or that deviated from Hardy-Weinberg equilibrium (*p* < 1×10^−3^). In addition, we required that variants should have all three genotypes (homozygous reference allele, homozygous alternative allele, and heterozygous genotypes) observed in the samples.

##### PCA of replication timing data

To account for potential batch effects and other unknown systematic biases in the replication timing data, we performed PCA using the filtered raw data with R function `prcomp`. Principal components (PCs) of the filtered raw data (“phenotype PCs”), along with the genotype PCs calculated above, were used as covariates in rtQTL mapping.

#### rtQTL mapping

##### Selection of phenotype PCs in rtQTL mapping

We followed the eQTL mapping framework used in the GTEx Project ^24^ (https://gtexportal.org/home/documentationPage) to map rtQTLs. We included the genotype (first three, similar to GTEx) and phenotype (first *k*) PCs in rtQTL mapping to account for non-genetic confounding factors. To find the optimal *k*, we tested each integer from 1 to 40. We consider the optimal *k* as the one leading to the highest number of windows harboring rtQTLs identified in rtQTL mapping. In this analysis, permutation parameter “`permut 50 500`” was used in fastQTL. Window level *p*-values were calculated, and the R package `qvalue` ^63^ was used to identify windows harboring rtQTLs at 10% FDR. This resulted in 24 and 22 selected as the optimal *k* for ESC and iPSC rtQTL mapping analysis, respectively, which was used in all subsequent rtQTL mapping analyses.

##### Cis*-rtQTL mapping using fastQTL*

We implemented a two-step approach to map rtQTLs using fastQTL ^64^. We generally restricted our analysis to *cis*-rtQTLs, defined as 1 Mb upstream or downstream of the center of each tested replication timing window. The first three genotype PCs and first 24 or 22 (for ESC and iPSC, respectively) phenotype PCs were included as covariates.

In the first step, we calculated a window-level *p*-value for each replication timing window using fastQTL, and then identified “significant windows”, i.e., windows with at least one significant rtQTL at 10% FDR, using the R package `qvalue`. This step is analogous to the identification of “eGenes” in eQTL mapping. For each window, fastQTL computes the lowest variant-level *p*-value and uses permutations to calculate the probability of observing a variant with equal or lower *p*-value under the scenario of no association, followed by beta approximation. Adaptive permutation parameter “`permut 1000 10000`” was used (similar to GTEx). We also repeated this step at 5% FDR.

In the second step, we identified genetic variants (referred to as SNPs for simplicity) associated with the “significant windows” identified in step 1, at 10% FDR. Here, we used a permutation-based strategy to determine the significance threshold for each tested window. By definition, FDR is the ratio of false positives (FP) to the sum of FP and true positives (TP). At a given *p*-value threshold *p*_*t*_, variants passing *p*_*t*_ are composed of both TP and FP. However, if we permute the phenotype, all variants with *p*-values lower than *p*_*t*_ are FP. Therefore, for a given window, FDR for a given *p*_*t*_ could be estimated as the mean number of variants passing *p*_*t*_ in permutations (i.e., all FP) divided by the number of variants passing *p*_*t*_ in the true association test (FP+TP). We then consider the maximum *p*_*t*_ with FDR ≤ 10% as the significance threshold of the window. The mean number of variants passing *p*_*t*_ in permutations was computed based on 500 permutations.

##### Evaluation of inflation of rtQTL mapping

To ensure that the computed variant-level *p*-values were not inflated, we calculated inflation factor with the Genomic Control method ^65^. We selected 200 windows (100 selected from windows carrying putative rtQTLs, and the other 100 randomly selected from the rest of the genome) and computed their association with genome-wide variants. We obtained variant-level statistics (which follows 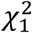 distribution under the null hypothesis) and computed the ratio of their median to the median of 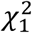 (0.456) as the genomic inflation factor. We calculated a genomic inflation factor (*λ*) as 1.03 and 1.00 for the ESC and iPSC dataset, respectively, thus the nominal *p*-values were not inflated; this was also supported by quantile-quantile plots.

##### Identification of rtQTLs

The following procedure was used to identify discrete rtQTLs, i.e., independent (not in LD) association signals, based on the significant SNPs mapped using the aforementioned two-step approach. For clarity, we denote independent association signals as rtQTLs, each of which contains multiple SNPs that are part of the association signal.

For each window, we identified all SNPs (if any) that passed the significance threshold. We selected the SNP with the lowest *p*-value as the “tag” variant of an rtQTL and assigned SNPs in LD (*r*^*2*^ ≥ 0.2) with the tag variant to the rtQTL. If there were any SNPs remaining that passed the significance threshold, we selected the SNP with the lowest *p*-value among the remaining SNPs as the tag variant of a new rtQTL and assigned all variants in LD with the new tag variant to the new rtQTL. This step was repeated until no variants passing the significance threshold were left. For the rtQTLs identified above, we kept only those with at least 10 variants and for which the *p*-value of the tag variant was less than 10^−3^.

For all calculations involving LD, data from the 1000 Genomes Phase 3 Europeans was used whenever available. For SNPs not called in the 1000 Genomes dataset, the current dataset was used for LD calculation.

Since nearby replication timing windows are highly correlated, the same rtQTL can be detected across multiple windows. We consolidated association signals detected in different windows if they satisfy all of the following three criteria: (1) the tag variants are in LD (*r*^*2*^ ≥ 0.2), (2) the replication timing windows are correlated (*R*^*2*^ ≥ 0.1), and (3) the distance between the windows is less than 2Mb.

In addition to separating rtQTLs by LD, we performed conditional association for each identified rtQTL. We conditioned on the top variant of each rtQTL and examined whether any SNPs that belong to this rtQTL still have significant association *p*-value (at *p* = 0.05 after Bonferroni correction). If so, this rtQTL was divided into multiple rtQTLs, each representing an independent association signal.

##### Filtering of rtQTLs

The putative rtQTLs identified were subjected to further filtering. First, we determined the boundaries of regions that significantly associated with each putative rtQTL. Starting at the window that most strongly associated with the tag variant (i.e., with the lowest *p*-value) of an rtQTL, we extended the region bi-directionally until the association was no longer significant (*p* > 0.05). We refer to this region as the “associated region”.

Next, we filtered false positives suspected to be potentially caused by short CNVs. During data processing (described above), we removed windows in which copy number measurement are potentially influenced by CNVs. However, short CNVs, spanning only one or two windows, may not have been detected and filtered and could lead to false positive rtQTLs (if they are in LD with SNPs). This type of false positive was identified by utilizing the raw unsmoothed data as follows: if a putative rtQTL is a false positive caused by a CNV, it would be (1) only observed in a small number of unsmoothed raw windows (overlapping with the CNV), and (2) will be more strongly associated with the raw data than with the smoothed data (in which the CNV will be smoothed within a broader region, thus decreasing association). Furthermore, it may have much stronger association with windows removed during replication timing data processing.

We computed the association *p*-values of the tag variant of each rtQTL with the (1) smoothed data within the associated region, (2) filtered raw data within the associated region, and (3) data that were removed during data processing within 1 Mb upstream or downstream of the associated region (referred to as “removed data” below).

Putative rtQTLs must satisfy all of the following criteria to be included in the final list of rtQTLs:

1. In the raw data, the tag variant must be associated (*p* < 0.05) with at least five windows.
2. The minimum *p*-value of the raw data must be higher (i.e., less significant), or no more than one order of magnitude lower, than that of the smoothed data.
3. The minimum *p*-value of the removed data must be higher, or no more than one order of magnitude lower, than that of the raw data. This criterion is relaxed to two or four orders of magnitude for rtQTLs with top *p*-value ≤ 5×10^−6^ and ≤ 5×10^−8^, respectively.
4. No more than two windows in the removed data have *p*-values lower than the minimum *p*-value for the raw data. This criterion is relaxed to three windows for rtQTLs with top *p*-value ≤ 5×10^−8^.
5. The minimum *p*-value from the raw data must be less than 0.01.
6. The associated region must be larger than one replication timing window.

In total, we identified 608 ESC rtQTLs, among which 603 were on autosomes and five were on the X chromosome in males. No rtQTLs were found on the X chromosome in females. This was not due to the reduced number of individuals tested, but likely resulted from the less structured replication timing profiles attributed to the female inactive X chromosomes: the similar-sized chromosome 7 had ten rtQTLs in the 50 male samples, not significantly different than the male X chromosome (*p* = 0.31, Fisher’s exact test), while there were fifteen rtQTLs on chromosome 7 in 66 female samples, significantly more than the none found on the female X chromosome (*p* = 7.41×10^−5^). We identified 1,167 iPSC rtQTLs. The nominal *p*-value of rtQTLs ranged from 1.02×10^−69^ to 9.63×10^−4^ (106 and 218 rtQTLs [17.4% and 18.7%] had *p* ≤ 5×10^−8^ in the ESC and iPSC dataset, respectively). The early- and late-replicating alleles were equally likely to be the reference allele (binomial *p* = 0.55), thus rtQTL mapping was not influenced by reference mappability bias.

##### Prioritizing causal genetic variants

For each rtQTL, CAVIAR^21^ was used to produce a shortlist of possible causal SNPs at 90% probability, from all SNPs in LD with the tag variant of the rtQTL (*r*^*2*^ ≥ 0.2). The shortlisted SNPs were used in all enrichment analyses.

##### rtQTL classification

We classified each rtQTL as affecting peak (initiation site), valley (terminus), or slope (transition region). For each rtQTL, we identified the replication timing loci that have large difference in replication timing (at least 90% of the maximum difference) between the early-replicating and late-replicating individuals (denoted as the “most variable replication timing loci”). We then calculated “fractional distance” of these loci along the peak-to-valley interval in which they reside. If a replication timing locus, with position *a*, resides in the interval between a peak (with position *b*) and a valley (with position *c*), its fractional distance was calculated as *a* minus *b*, divided by *c* minus *b*. We considered an rtQTL as affecting an initiation site if the fractional distance of at least one of the most variable replication timing loci was less than 0.3. Conversely, we considered an rtQTL as affecting a valley if the fractional distances of all of the most variable replication timing loci were greater than 0.7. rtQTLs that did not fall into either of these two categories were categorized as affecting slopes.

We further classified rtQTLs that affect peaks based on whether the top rtQTL SNP was located proximal or distal to the peak. Specifically, we calculated fractional distance of the top rtQTL SNP for each rtQTL that affect peaks, using the same approach as described above. The top rtQTL SNP was considered proximal to the peak if its fractional distance was less than 0.3 and was considered distal to the peak otherwise.

##### Merging ESC and iPSC rtQTLs

We combined ESC and iPSC rtQTLs for a number of analyses. To minimize double counting of rtQTLs discovered in both datasets, we generated a merged rtQTL list for these analyses. This list excluded iPSC rtQTLs that met the following criteria: (1) a genetic variant that belongs to the given iPSC rtQTL and has a *p*-value no more than two orders of magnitute higher than the top *p*-value of the iPSC rtQTL also belongs to a ESC rtQTL, and (2) the direction of effect of the given genetic variant is the same in the iPSC and ESC datasets. We merged the 608 ESC rtQTLs and 1,167 iPSC rtQTLs into a list of 1,617 combined rtQTLs.

#### Validation of rtQTLs

To validate the iPSC rtQTLs, we examined their reproducibility in the Merkle_batch1 ESC dataset (108 European ancestry samples only). Validation was performed using fastQTL^64^ by testing the association between the strongest rtQTL SNP and the replication timing locus closest to the locus with the strongest association in the discovery set (HipSci iPSCs). Three genotype PCs and 24 phenotype PCs were included as covariates. When the strongest rtQTL SNP was not available in the validation dataset (Merkle_batch1 ESCs), an rtQTL SNP from the same rtQTL that has *p*-value less than two orders of magnitude higher than that of the strongest rtQTL SNP was used instead. We found that the -log_10_(*p*-values) of rtQTLs are highly correlated between the discovery and validation datasets (*r* = 0.75, *p* = 1.28×10^−176^). We then repeated this analysis in the opposite direction (validate ESC rtQTLs using HipSci iPSCs) and obtained similar results (*r* = 0.76, *p* = 7.81×10^−113^). These observations support that the rtQTLs identified in this study are highly reproducible.

We also used three additional datasets to validate ESC rtQTLs. The first dataset contains 9 hESCs in Merkle_batch2 and the 8 hESCs in Merkle_batch1 that were excluded in rtQTL mapping due to ancestry. The second and third datasets are the in-house hESC and iPSC dataset, respectively.

For the first dataset, validation was performed in fastQTL. Validation using the second and third datasets were performed in MATLAB by calculating the Pearson correlation *p*-value between the strongest rtQTL genetic variant and the replication timing locus with the strongest association in the discovery set. We tested rtQTLs of which the top genetic variant was polymorphic and had all three genotypes in the validation dataset. rtQTLs were excluded if the alternative allele of the top genetic variant in the validation dataset was not consistent with that of in the discovery set. This left 427 regions that could be tested in the third dataset, and 396 regions in the fourth dataset. Replication timing of these two datasets were inferred using GenomeSTRiP (as described above) in 2.5Kb windows of uniquely alignable sequence^59^. For each sample, windows with copy number >3 or <1 were removed. We used a segmentation algorithm (`segment` in MATLAB) to further remove outlier data points (segments with mean >2.45 or <1.55 were removed). The data was then smoothed using cubic smoothing spline with parameter 10^−17^.

We considered an rtQTL as “validated” if it was associated with replication timing with nominal *p* < 0.05 and had the same direction of effect in at least one of the validation datasets. The binomial test was used to assess significance of the number of validated rtQTLs, with binomial parameter calculated as 1–(1–0.05/2)^4^ = 0.0963 (i.e., the probability under random chance that an rtQTL will be validated in at least one dataset).

##### SMARD

SMARD analysis was carried out as previously described^22^. Briefly, cells were pulse labeled sequentially with 25 μM IdU and CldU. The cells were then embedded in 1% InCert agarose and lysed. The remaining embedded genomic DNA was digested with restriction endonucleases. Pulsed field gel electrophoresis (PFGE) was used to separate DNA according to size. The segment containing the locus-of-interest was identified with Southern blot and the gel slice was excised. Agarose was then melted, and individual DNA strands were stretched on silanized glass slides. Immunostaining was employed to detect the halogenated nucleotides in the replicated DNA. Biotinylated FISH probes were used to identify DNA molecules containing the locus-of-interest.

#### Multi-rtQTLs

To identify multi-rtQTL regions, we considered separate rtQTLs to be associated with the same region if the replication timing loci most strongly associated with them were correlated (*R*^*2*^ ≥ 0.2) across individuals, were in physical proximity (< 2 Mb apart), and each provided additional explanatory power for replication timing. Secondary rtQTLs were either not in LD with the primary ones (130 and 265 multi-rtQTL regions in the ESC and iPSC dataset, respectively), or provided additional explanatory power despite being in LD (5 cases in ESC and 10 cases in iPSC).

Some analyses were performed with ESC and iPSC multi-rtQTL regions combined. To avoid double-counting in these analyses, we excluded iPSC multi-rtQTL regions that has at least one rtQTL that was also found in the ESC dataset. We combined 135 ESC and 275 iPSC multi-rtQTL regions into 318 multi-rtQTL regions.

We examined the possible interaction between primary and secondary rtQTLs in regions with two, three, and four rtQTLs. If there was no interaction, we expect that the replication timing in these regions will be positively linearly correlated with the dosage of early-replicating alleles. To enable pooling of multi-rtQTL regions for Fig. 2E and 2F, we normalized replication timing for the loci with strongest association with the primary rtQTL of each multi-rtQTL region to *Z*-score (by subtracting the mean and dividing by the standard deviation of replication timing of this locus among samples) and denoted them as relative replication timing. They were pooled and linear regression analysis was performed using the R `lm` function.

We used a likelihood-ratio test to assess whether the additive or synergistic models better explained replication timing at multi-rtQTL regions. We tested the null hypothesis by which replication timing is proportional to the number of early-replicating rtQTL alleles carried by an individual at a multi-rtQTL region (additive effect), against the alternative, by which replication timing is more extremely biased in individuals carrying multiple early (or late) rtQTL alleles (synergistic interaction). We used 58 regions that harbored two rtQTLs and had at least one individual with zero and one with four early-replicating alleles. We fitted two linear models, with the response variable being replication timing and explanatory variable being genotype dosage. In the null (additive) model, genotype dosage was between zero to four, matching the number of early-replicating alleles that individual carried. In the alternative (synergistic) model, genotype dosages of individuals carrying zero or four early-replicating alleles were estimated from actual data. We then compared −2×(log likelihood ratio) with the chi-squared distribution with two degrees of freedom to obtain a *p*-value.

We examined whether the primary and secondary rtQTLs in ESC were in close spatial proximity in nuclear space. We obtained Hi-C contact matrix of the H1 cell line from Juicebox^66^ and computed contact score between each pair of primary and secondary rtQTLs. We compared the median of these scores with 100 permutations, in which the distances between primary and secondary rtQTLs were preserved but actual genomic locations were randomly shifted between 1 and 2 Mb up- or downstream. *P*-value was computed using *Z* score, with mean and standard deviation estimated from the permutations.

#### Epigenetic enrichment analyses

##### Data sources

Chromatin state and histone mark data for eight human ESC lines (seven of which are included in our primary replication timing data) and five human iPSC lines were obtained from the Roadmap Epigenomics Project^39^. For the analyses of overall enrichment of epigenetic features at rtQTL locations, we combined (i.e., took the union of) histone peaks and chromatin state calls from the eight cell lines. For histone marks, observed data was used when available, and imputed data (from ChromImpute^40^) was used when observed data was not available. Imputed data were used for plotting of histone tracks. Binding site information for 51 TFs was obtained from the ENCODE Project^43^. SOX2 binding site information was obtained^46^. TFs with binding sites that overlapped < 15 rtQTLs were excluded from this analysis.

##### Enrichment calculations

For each feature (chromatin state, histone marks, TF, etc.), we are interested in the number of rtQTLs that have at least one SNP overlapping with the feature, and whether this is more or less likely (i.e., enriched or depleted) than expected by chance. Statistical significance was assessed with one-tailed binomial test. The binomial parameter *p* was estimated from 100 random permutations, from which we estimated the probability of random SNPs (matched with the rtQTLs, see details below) overlapping with the feature. Correction for multiple testing was applied when multiple features from the same category (e.g., histone marks) were tested.

For each rtQTL, we searched for random SNPs that match the characteristics of the tag variant of the rtQTL (denoted as “actual tag variant”) and used the matched variants (“matched tag variants”) to tag the random sets of SNPs used in permutations. We required that the matched tag variants must be at least 2 Mb away from the actual tag variant. The matched tag variants must also have satisfied all three following criteria: (1) have similar minor allele frequency (< 5% difference), (2) have similar distances to the nearest replication initiation site and terminus (< 50 kb difference), and (3) have similar replication timing (< 0.5 units difference) with the actual tag variant. We require the matched tag variants to have the same number, or more, SNPs in LD (*r*^*2*^ ≥ 0.2) than the actual tag variant.

In each permutation, and for each rtQTL, we constructed a set of random SNPs using SNPs in LD with a randomly selected matched tag variant. The number of variants in the set is the same number of variants included in the actual rtQTL. Eleven (1.82%) rtQTLs in hESC and 41 (3.51%) rtQTLs in iPSC that had less than 200 matched tag variants genome-wide were excluded from the analysis.

Enrichment analyses were also repeated using epigenetic data from one hESC line only (as opposed to combining data from eight hESC lines), with consistent results.

#### Using epigenetic features to predict replication initiation site locations

##### Identification of epigenetic feature combinations

To identify combinations of chromatin marks enriched at rtQTLs, we used a stepwise, iterative approach. The hESC rtQTLs and epigenetic data were used. We considered 29 histone marks (Fig. S2C) and also included H2A.Z, DNase I hypersensitivity, and binding sites of 51 TFs and other DNA binding factors (referred to as TFs for simplicity).

First, we tested each individual epigenetic feature (histone mark or TF) to identify features that are enriched at rtQTL SNPs. Enrichment was examined using the same permutation-based approach described above. The only difference was that each rtQTL individual SNP was considered independently (as opposed to being considered together with other SNPs assigned to the same rtQTL), as our goal was to identify co-occurrence of epigenetic features at the same exact genomic locations. Statistical significance was assessed using Fisher’s exact test. We corrected for multiple testing at 5% FDR using the Benjamini-Hochberg correction.

Next, for each enriched feature identified in the first step, we examined whether the pairwise combination of this feature and any of the other epigenetic features has stronger enrichment. Specifically, we restricted the enrichment analysis to the rtQTL SNPs that carry the enriched feature and tested whether the additional epigenetic feature is enriched in the set of restricted rtQTL SNPs. This step was repeated iteratively, each round restricting the analysis to the enriched combinations of epigenetic features identified in the previous round, until no further enrichment was found. In Fig. 3A, combinations containing TFs were not included for simplicity and since they were not carried through to the four- and five-mark combinations.

To identify “me^3^ac^hyper^” regions, we first identified regions that carry one of the 13 five-mark combinations and kept regions that overlap with peaks from at least 11 variable acetylation marks. We merged me^3^ac^hyper^ regions that co-occurred within 10 kb. In Fig. S4A–C, the position of initiation sites found in >10% of the samples were determined based on local maxima in the averaged replication timing profile. When calculating distances (fractional and physical), distance was set to zero for me^3^ac^hyper^ regions that overlap with an initiation site (i.e., the interval between boundaries of the initiation sites). If a me^3^ac^hyper^ region does not overlap with any initiation site, its physical distance was calculated as the distance to the nearest initiation site boundary. To explore the independence of the replication initiation histone code from gene expression, we divided me^3^ac^hyper^ regions into two classes, based on whether there were TSS of expressed gene(s) (mean RPKM in ES cell lines > 0.5) within a given me^3^ac^hyper^ region. RNA-seq data was obtained from the Roadmap Epigenomics Project^39^. We then compared the positive predictive value for these two classes of me^3^ac^hyper^ regions.

##### Receiver Operating Characteristic (ROC) curves

To obtain ROC curves, we used various thresholds (see below) to predict whether a replication timing window corresponds to a replication initiation site. Specifically, we predict a window as being an initiation site if it was located within *k* kb of a me^3^ac^hyper^ region (in hESCs and iPSCs) or a region that carries the H3K4me3-H3K9me3-H3K36me3 combination (in LCLs). We used various values for *k*, from 0 to 2,000. We then compared the prediction with actual data (whether the replication timing windows fell within the boundaries of the identified initiation sites) to calculate true and false positive rates. For permutations, we randomly shifted the locations of the me^3^ac^hyper^ regions between 1 Mb and 2Mb and obtained ROC curves and AUC_ROC_ based on these random intervals.

##### Replication initiation site prediction in LCLs and iPSCs

We assessed the generalizability of the replication initiation histone code in LCLs and iPSCs. LCL is a cell type that has a distinct epigenetic and replication landscape from hESC lines^23,67,68^, and iPSCs have similar but not identical to replication timing profiles to hESCs (*r =* 0.90). Replication timing profile for the GM12878 LCL and 192 unrelated iPSCs were inferred from whole-genome sequencing data^19,45^. For iPSCs, initiation site locations were identified based on the averaged iPSC replication timing profile. When calculating physical distance of predicted initiation sites to actual initiation sites, we defined initiation site boundaries as 100 kb upstream and downstream of the local maxima in the replication timing profiles. Data for H3K4me3, H3K9me3, and H3K36me3 for the GM12878 LCL was from the ENCODE Project^43^. Additional data of H3K4me3 and H3K36me3 for 18 LCLs was obtained from^44^, and merged with the ENCODE data. Histone mark data for five iPSCs was from the Roadmap Epigenomics Project^39^. If a histone code location was found in one cell type (either hESC or LCL), but no histone code location was found within 100 kb in the other cell type, we denoted this region as cell-type-specific. Otherwise, this region was denoted as “shared” between the two cell types.

#### Identification of features associated with replication timing

##### Chromatin states and histone marks

Replication timing data was available for seven of the eight hESC lines that were analyzed in the Roadmap Epigenomics Project. Using rtQTL and epigenetic data from these seven cell lines, we designed an analysis to identify chromatin states and histone marks associated with replication timing. The rationale is that epigenetic features promoting earlier replication would be more likely to be carried by early-replicating-associated rtQTL genotypes, and *vice versa* for late replication. We were only able to perform this analysis for hESC rtQTLs because we did not have replication timing, genotype, and epigenetic data for the same iPSC lines.

We aggregated information from all rtQTL SNPs, except those that are monomorphic in the seven cell lines. We assigned each cell line by genotype to one of three categories, i.e., early-replicating, heterozygous, and late-replicating, at each rtQTL SNP. For each epigenetic feature, we tested whether the cell lines carrying the early-replicating genotypes are more (or less) likely to harbor it than the cell lines carrying the late-replicating genotypes, using the two-tailed binomial test. The binomial parameter *p* was calculated as *p*_*late*_ × (*p*_*perm_early*_ / *p*_*perm_late*_), where *p*_*late*_ is the proportion of late-replicating genotypes carrying this feature, and *p*_*perm_early*_ and *p*_*perm_late*_ are estimated from ten permutations (described below). Bonferroni correction was used to correct for multiple testing at the 0.05 level.

In each permutation, we used random SNPs matched for rtQTLs (for details see the “enrichment analyses” section), and randomly designated one genotype as the early-replicating genotype. We obtained genotype and epigenetic information from the seven cell lines at these random SNPs and calculated the proportion of early-and late-replicating genotypes carrying the feature in ten permutations (*p*_*perm_early*_ and *p*_*perm_late*_).

We examined the relationship between early-replicating genotypes and expression of nearby genes (within 200 kb of rtQTLs). Array-based expression data was obtained for nine ES cell lines^69^ for which replication timing data was also available. Genes with mean expression level > 1 were used, and expression level was normalized within each gene. We aggregated the expression levels of all genes near all rtQTLs for the nine ES cell lines (except for rtQTLs that were monomorphic in these cell lines), and tested the correlation between expression level and the number of early-replicating alleles.

#### Transcription factors

To identify TFs that regulate replication timing, we tested whether rtQTL alleles (in the CAVIAR 90% causal set) influence the binding affinity (motif score) of 21 TFs^70^. Under the hypothesis that some rtQTLs function by altering sequence motifs of TFs that promote or repress replication, early-replicating alleles will be more likely to have higher binding affinities than late-replicating alleles to the TFs that promote earlier replication, and *vice versa* for late-replicating alleles. We used this principle to identify TFs associated with replication timing. We tested the motifs of all TFs studied in Fig. S5A, if available. Of note, SOX2 was not included in this analysis because its motif information was not available. This analysis was repeated with iPSC rtQTLs. We were not able to perform the analysis described above for chromatin states and histone marks with TFs because TF ChIP-seq data was only available for one hESC or iPSC line.

TF binding affinity data, measured by motif score, was obtained from HaploReg^70^. Sequence logos for TF binding motifs were created using WebLogo 3^71^. For each rtQTL SNP, motif scores of the two alleles were obtained for the TFs, and their difference is the log_2_ fold difference in probability that the allele is in a binding motif of the given TF. Higher difference in motif scores means that this SNP can more substantially alter the binding affinity of this TF.

For each TF, we counted the weighted number of rtQTLs for which the early-replicating (or the late-replicating) allele had higher predicted binding affinity, weighted by the difference in motif scores between the two alleles, i.e., rtQTLs with a higher motif score difference will have heavier weight. This weighting scheme assigns heavier weights to those rtQTLs for which the changes in allele state will result in more substantial change in TF binding affinity. If there were more than one potential causal SNPs of an rtQTL located within binding motifs of a given TF, the SNP with the lowest *p*-value was used. We compared the numbers to permutations, in which SNPs matched for rtQTLs were randomly selected and the early-replicating alleles were randomly assigned, using the chi-squared test for a 2×2 contingency table. This test assesses whether the early-replicating alleles are more (or less) likely to have higher TF binding affinity than late-replicating alleles than expected by chance. Benjamini-Hochberg correction at 10% FDR was used to correct for multiple testing.

For OCT4, NANOG, and CTCF (for which there are abundant ChIP-seq data available in hESC), we repeated this analysis using only motifs that overlap with TF ChIP-seq peaks (i.e., confirmed TF binding). Consistent with the results in Fig. 6A, we found that OCT4 and NANOG were significantly more likely to bind early-replicating alleles (*p* = 5.97×10^−7^ and 2.62×10^−15^; log_2_ ratio improved from 0.27 and 0.29 to infinity and 2.58, respectively), while CTCF was significantly more likely to bind late-replicating alleles (*p* = 0.02, log_2_ ratio improved from −0.22 to −1.19).

Supplementary Figures were included in the main text near where they were mentioned.

Table S1. List of rtQTLs Identified in This Study.

For the last column (“classification”), “VALLEY” or “SLOPE” denotes that this rtQTL affects valley or slope, respectively. “PEAK (SNP proximal)” or “PEAK (SNP distal)” denotes that this rtQTL affects peak, and the top rtQTL SNP is proximal or distal to the peak, respectively.

This table is provided in a separate Excel spreadsheet.

